# Enhancer priming enables fast and sustained transcriptional responses to Notch signaling

**DOI:** 10.1101/497651

**Authors:** Julia Falo-Sanjuan, Nicholas C Lammers, Hernan G Garcia, Sarah Bray

## Abstract

Information from developmental signaling pathways must be accurately decoded to generate transcriptional outcomes. In the case of Notch, the intracellular domain (NICD) transduces the signal directly to the nucleus. How enhancers decipher NICD in the real time of developmental decisions is not known. Using the MS2/MCP system to visualize nascent transcripts in single cells in *Drosophila* embryos we reveal how two target enhancers read Notch activity to produce synchronized and sustained profiles of transcription. By manipulating the levels of NICD and altering specific motifs within the enhancers we uncover two key principles. First, increased NICD levels alter transcription by increasing duration rather than frequency of transcriptional bursts. Second, priming of enhancers by tissue-specific transcription factors is required for NICD to confer synchronized and sustained activity; in their absence, transcription is stochastic and bursty. The dynamic response of an individual enhancer to NICD thus differs depending on the cellular context.

## Introduction

Genes respond to external and internal cues through the action in the nucleus of transcription factors and effectors of signalling pathways. Regulatory regions that surround genes, termed enhancers, integrate the information from these inputs to produce an appropriate transcriptional output. During development some of these decisions occur in a matter of minutes, but usually the outcomes are measured many hours later. Rarely have transcription dynamics been analyzed *in vivo* in the real-time of developmental signalling pathways, so we know little about how recipient enhancers decipher the signals. For example, enhancers could respond in a digital manner, working as simple on-off switches, or as analog devices, operating as a rheostat so that signal levels can modulate the output (Blackwood and Kadonaga 1998; Garcia et al. 2013; Lammers et al. 2018). In either case they must also have the capability to detect and transduce key parameters to the transcription machinery, such as signal duration and thresholds.

With the advent of precise and quantitative methods to measure transcription, such as single molecule fluorescence in situ hybridization (smFISH) or live imaging, it has become evident that transcription is not a continuous process. Instead, genes actively transcribed undergo bursts of initiation that are often separated by inactive intervals (Chubb et al. 2006; Golding et al. 2005). Bursting is thought to occur because the dynamics of enhancer-promoter activation leads to episodic polymerase release. One consequence of this is that factors modulating the levels of transcription can do so by changing either the frequency with which a burst occurs (measured by the gap between bursts) or the size of each burst (measured by changes in burst duration and/or amplitude). Since forced looping of the *beta-globin* enhancer to its promoter led to an increase in burst frequency (Bartman et al. 2016), it has been proposed that transcription factors activate transcription by modulating enhancer-promoter interactions, and hence bursting frequency; although other studies suggest enhancer-promoter interactions are not the underlying basis of transcriptional bursting (Lim et al. 2018; Chen et al. 2018). Though the molecular origin of bursting remains unknown, bursting frequency rather than burst duration or amplitudes seems to be the major parameter modulated in different species and contexts (So et al. 2011; Senecal et al. 2014; Xu et al. 2015; Desponds et al. 2016; Padovan-Merhar et al. 2015; Lammers et al. 2018; Berrocal et al. 2018). For example, enhancers controlling early patterning genes in *Drosophila* embryos produce similar bursting size but have different bursting frequencies, which can be attenuated by the presence of insulators (Fukaya et al. 2016). Similarly, steroids increase the bursting frequency of target enhancers (Larson et al. 2013; Fritzsch et al. 2018). However, it remains to be discovered whether all transcription factors alter transcription dynamics in this way and specifically whether it is these or other properties that are modulated by developmental signals to confer appropriate outputs in the *in vivo* setting of a developing organism.

Transcriptional bursting is thought to make an important contribution to heterogeneity in transcriptional activity between cells (Raj and Oudenaarden 2008). For example, in cells exposed to estrogen, response times for transcription activation were highly variable and there was no coherent cycling between active and inactive states (Fritzsch et al. 2018). Stochastic transcriptional behaviour is of key importance in several developmental decisions, such as differentiation of photoreceptors in the *Drosophila* eye (Wernet et al. 2006), hematopoietic cell differentiation in mouse cells (Chang et al. 2008; Ng et al. 2018) or during neuronal differentiation in the zebrafish retina (Boije et al. 2015). But while an attractive feature for promoting heterogeneity, inherent variability in responses could be extremely disruptive in developmental processes where the coordinated response of many cells is required to pattern specific structures. In some cases this may be circumvented by averaging mechanisms that allow cells to produce homogeneous patterns of gene expression (Little et al. 2013). For example, cells that express the mesodermal determinant Snail average their transcriptional output over a period of 20 minutes by mRNA diffusion to produce a homogeneous field of cells and a sharp boundary in *Drosophila* syncytial embryos (Bothma et al. 2018). However it is only in rare circumstances that mRNA diffusion can operate and it is unclear whether other averaging mechanisms would be effective over shorter time intervals. To effectively achieve reproducible patterns, cells must therefore overcome the variability that is inherent in transcriptional bursting and stochastic enhancer activation.

Notch signaling is a highly conserved developmental signaling pathway that is deployed in multiple contexts. It has the unusual feature that the Notch intracellular domain (NICD) transduces the signal directly to the nucleus, when it is released by a series of proteolytic cleavages precipitated by interactions with the ligands. NICD then stimulates transcription by forming a complex with the DNA binding protein CSL and the co-activator Mastermind (Mam) (Bray 2006). The lack of intermediate signalling steps and amplification makes this a powerful system to investigate how signals are deciphered by responding enhancers. Furthermore, there may be differences in the levels and dynamics of NICD produced by different ligands (Nandagopal et al. 2018). However, although its role as a transcriptional activator is well established, at present we know little about how enhancers respond to NICD in the real time of developmental decisions. For example, do enhancers operate as simple switches, detecting when NICD crosses a threshold? Or are they sensitive to different levels of NICD, in which case does NICD, like other factors, modulate bursting frequency? Nor do we know what features in the sequence of the responding enhancers confer the output properties, although it has been suggested that enhancers with paired CSL motifs (referred to as SPS motifs) (Bailey and Posakony 1995; Nam et al. 2007), whose precise spacing could favour NICD dimerization, have the potential to yield the strongest responses (Nam et al. 2007).

In order to determine how enhancers respond to Notch activity in real time we have used the MS2/MCP system to visualize nascent transcripts in *Drosophila* embryos. To do so we used two well-characterised Notch responsive enhancers that drive expression in a stripe of mesectoderm (MSE) cells and analyzed their transcription profile over time at the single cell level. Strikingly all MSE cells initiate transcription within a few minutes of one another, and once active, each nucleus produced a sustained profile of transcription. By manipulating the levels of NICD and altering key motifs within the enhancers we uncover two key principles. First, the ability of NICD to confer synchronized and sustained activity in MSE requires that the enhancers are primed by tissue-specific transcription factors. In their absence, MSE enhancers confer stochastic bursty transcription profiles, demonstrating that different response profiles can be generated from a single enhancer according to which other factors are present. Second, changing Notch levels modulates the transcription burst size but not length of the periods between bursts, in contrast to most current examples of enhancer activation. These two key concepts are likely to be of general importance for gene regulation by other signalling pathways in developmental and disease contexts.

## Results

### Synchronised and sustained enhancer activation in response to Notch

To investigate how Notch signals are read out by an enhancer in real time, we focused on well-characterized mesectodermal (MSE) enhancers from the *Enhancer of split-Complex (E(spl)-C)* (known as *m5/m8*) and from *singleminded* (*sim*) (Morel and Schweisguth 2000; Cowden and Levine 2002; Zinzen et al. 2006a). These direct expression in two stripes of MSE cells during nuclear cycle 14 (nc14) when Notch is activated in response to Delta signals from the presumptive mesoderm (Fig. 1**AB**) (Morel et al. 2003; De Renzis et al. 2006; Zinzen et al. 2006a). The MSE converges to the midline during gastrulation, ultimately forming CNS midline precursors similar to the vertebrate floorplate. To visualize transcription from the MSE enhancers in real time, and define the response properties conferred by a defined enhancer DNA sequence, they were inserted into MS2 reporter constructs containing the promoter from *even-skipped* (*peve*), 24 *MS2* loops and the *lacZ* transcript (Fig. 1**A**). When these *MS2* reporters were combined with MCP-GFP in the same embryos, nascent transcription was marked by the accumulation of MCP-GFP in bright nuclear puncta, where the total fluorescence in each spot is directly proportional to the number of transcribing mRNAs at any timepoint (Fig. 1**AB**)(Garcia et al. 2013). In this way the levels of transcription can be followed over time at the single cell level by tracking the puncta relative to nuclei (labelled with His2Av-RFP).

**Figure 1.**
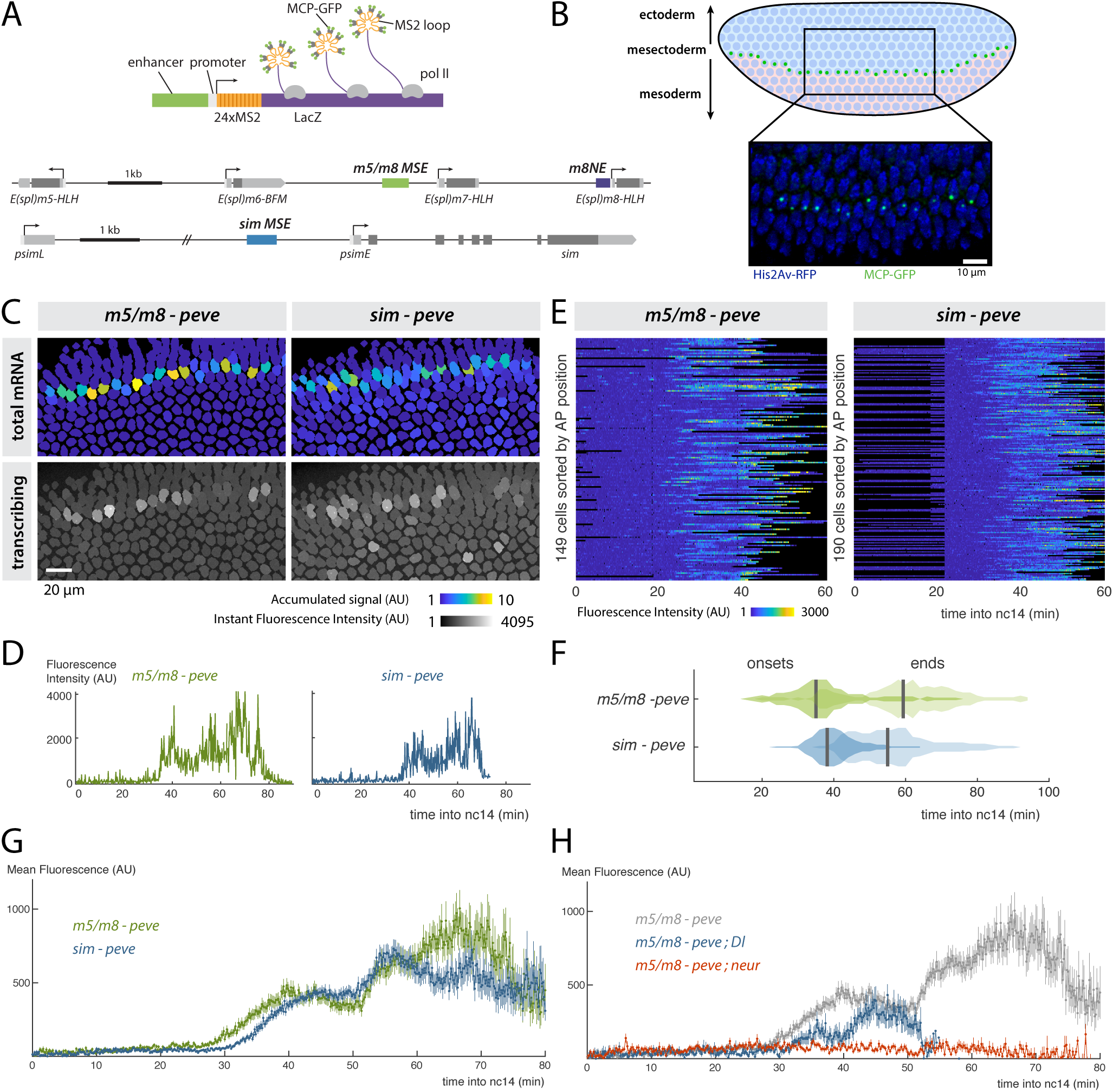
Synchronous activity of two Notch responsive enhancers. **A**) Diagrams illustrating the strategy for live imaging of transcription using the MS2 system (top) and the location of mesectoderm (MSE) and neuroectoderm (NE) enhancers in *E(spl)-C* (*m5/m8*; *m8NE*) and *single minded* (*sim*) (bottom). Arrows indicate transcription start-sites, boxes in lower panel indicate promoters (white), non-coding (light grey) and coding (dark grey) transcribed regions. **B**) Diagram of a blastoderm *Drosophila* embryo, indicating Delta expression (pink) in the mesoderm which activates Notch in a flanking stripe of cells (green dots) to specify the MSE. Image: transcription from *m5/m8* detected by accumulation of MCP-GFP in bright puncta (green), nuclei are labelled by His2Av-RFP (blue). **C**) Tracked expression from *m5/m8* and *sim* reporters. Top panels: tracked nuclei false-colored by total signal levels, proportional to total mRNA production. Bottom panels: single frame of *m5/m8* and *sim* embryos, tracked nuclei shaded according to maximum pixel intensity in that frame. **D**) Examples of *m5/m8* and *sim* fluorescence profiles from individual nuclei that exhibit ‘sustained’ activity. **E**) Heatmaps representing fluorescence profiles of *m5/m8* and *sim* in all MSE nuclei during nc14 (scale as indicated with blue, no expression, yellow, high expression; black indicates periods where nuclei were not tracked). **F**) Distributions of onsets and end-points of transcription from *m5/m8* and *sim* in the MSE. **G**) *m5/m8* and *sim* produce similar average temporal profiles. Mean fluorescent intensity of MCP-GFP puncta at the indicated times after start of nc14. **H**) Transcription from *m5/m8* is curtailed in embryos lacking zygotic production of Delta (Dl) and abolished in embryos lacking *neuralized* (neur). Grey trace, profile from *m5/m8* in wild-type embryos shown in **G**. In **G** and **H** mean and SEM of all MSE cells are shown. n = 3 (*m5/m8*), 3 (*sim*), 2 (*m5/m8; Dl*), 2 (*m5/m8; neur*) embryos.

By visualizing transcription in real time we could see that *m5/m8* and *sim* were both activated in all MSE cells within a narrow time-window (~ 10 min) in nc14 (Fig. 1**CEF**, **Movie S1**, **Movie S2**). Activity was then maintained in these nuclei throughout the remaining period of nc14 as the embryos underwent gastrulation. They exhibited what we refer to as “sustained activity” because each puncta retained high levels of fluorescence rather than exhibiting clearly distinct bursts (Fig. 1**D**), although we note that the resolution of bursting events is limited by the time each polymerase takes to complete transcription (estimated as 1.6-2.5 min for these reporters) (Fukaya et al. 2017). Transcription then ceased after 30-50 minutes, with less synchrony than at the onset (Fig. 1**F**). Identical response profiles were obtained when the *m5/m8* reporter was inserted at a different genetic locus (Fig. S1**AB**).

Sustained activity is a feature of *m5/m8* and *sim* and not a general property of Notch responsive enhancers at this stage, as a neurectodermal enhancer from *E(spl)m8-bHLH* (*m8NE*, Fig. 1**A**) produced profiles where individual bursts of activity were clearly resolved, which we refer to here as ‘bursty’ (Fig. S1**ABC**). Furthermore, even though the profiles produced by *m5/m8* and *sim* were continuous, the amplitude fluctuated, likely reflecting episodic polymerase release. However it is notable that *m5/m8* and *sim* both directed transcription profiles that were highly co-ordinated temporally, with each conferring a prolonged period of activity that was initiated within a short time-window (Fig. 1**EF**). Indeed, the mean profile of all MSE cells analyzed was almost identical for the two enhancers (Fig. 1**G**). This is remarkable given that they contain different configurations of binding motifs and implies that the mesectoderm cells undergo a highly synchronized period and level of Notch signaling.

To assess the relative contributions of the enhancer and promoter to response profiles, we next tested the consequences from substituting different promoters with *m5/m8* and *sim* enhancers, inserting the reporters at the same genomic position to ensure comparability. First, when *peve* was replaced by a promoter from *sim* (*psimE*), both *m5/m8* and *sim* produced lower levels of transcription, but the overall temporal profiles and mean levels remained similar (Fig. S1**D**). Second, when we combined *m5/m8* with another heterologous promoter, *hsp70*, or with four promoters that could be interacting with *m5/m8* in the endogenous *E(spl)-C* locus, they similarly led to changes in the mean levels of transcription without affecting the overall temporal profile or expression pattern (Fig. S1**E**). Notably, even in combinations yielding lower levels, the transcription profiles remained sustained rather than breaking down into discrete bursts (Fig. S1**F**) consistent with promoters affecting mean levels of activity without modulating bursting frequencies. Of those tested, *pm6* produced the lowest mean levels (Fig. S1**E**), which correlates with the inactivity of *E(spl)m6-BFM* at this stage and argues for an underlying enhancer-promoter compatibility at the sequence level (Fig. S1**E**) (Zabidi et al. 2014). However, there was no obvious relationship between the mean levels of transcription produced by a promoter and the presence or absence of sequence motifs for factors associated with promoter accessibility, such as Zelda or Trithorax-like (Blythe and Wieschaus 2016). Nor was there a correlation between their activity levels with *m5/m8* in the MSE and those with a heterologous developmental enhancer in *Drosophila* S2 cells (Arnold et al. 2016). Nevertheless, the fact that similar temporal profiles were produced with all the promoters confirms that the enhancers are the primary detectors of Notch signaling activity.

To verify that MSE transcription was Notch-dependent we measured transcription from *m5/m8* in embryos where Notch activity was disrupted by mutations. Embryos lacking Neuralized, an E3 ubiquitin ligase required for Delta endocytosis that is critical for Notch signalling (Morel et al. 2003; De Renzis et al. 2006), had no detectable transcription from *m5/m8* in the MSE (Fig. 1**H**). Likewise, *m5/m8* activity was severely compromised in embryos carrying mutations in *Delta*. Because Delta protein is deposited in the egg maternally (Kopczynski et al. 1988), these embryos contained some residual Delta which was sufficient for a few scattered MSE cells to initiate transcription (Fig. S1**G**). However their transcription ceased prematurely, within <20 min (Fig. 1**H**, S1**G**). Together these results confirm that the enhancers require Notch signalling for their activity in the MSE, in agreement with previous studies (Morel and Schweisguth 2000; Zinzen et al. 2006a), and further show that continued Notch signalling is needed to maintain transcription, arguing that the MSE enhancers also detect persistence of NICD.

### Coordinated activity of enhancers within each nucleus

Although *m5/m8* and *sim* confer well coordinated temporal profiles of transcriptional activity, there is nevertheless cell to cell variability in their precise time of activation (Fig. 1**F**). To investigate whether this cell to cell variability was due to the stochastic nature of transcription (intrinsic variability) or whether it indeed reflects differences in signalling from Notch (extrinsic variability) (Elowitz 2002; Raser and Shea 2006) we monitored expression from two identical alleles of the MS2 reporters (Fig. 2**A**). Transcription from these two physically unlinked loci were detected as distinct puncta in each nucleus so that they could be tracked independently. We found a remarkable synchrony in the onset of transcription from both alleles of a given enhancer (Fig. 2**B**). More than 80% of the cells initiated transcription from both alleles within 5 min, (Fig. S2**C**). This contributes to ~ 6-30% of the total variability (Fig. 2**D**), indicating that most of the onset variability was due to extrinsic factors. There was less synchrony in the time at which transcription was extinguished (Fig. 2**B** S2**A**), but the extent of intrinsec variability was still much lower than that between cells (only contributing to less than 15% of the total variability, Fig. 2**D**).

**Figure 2.**
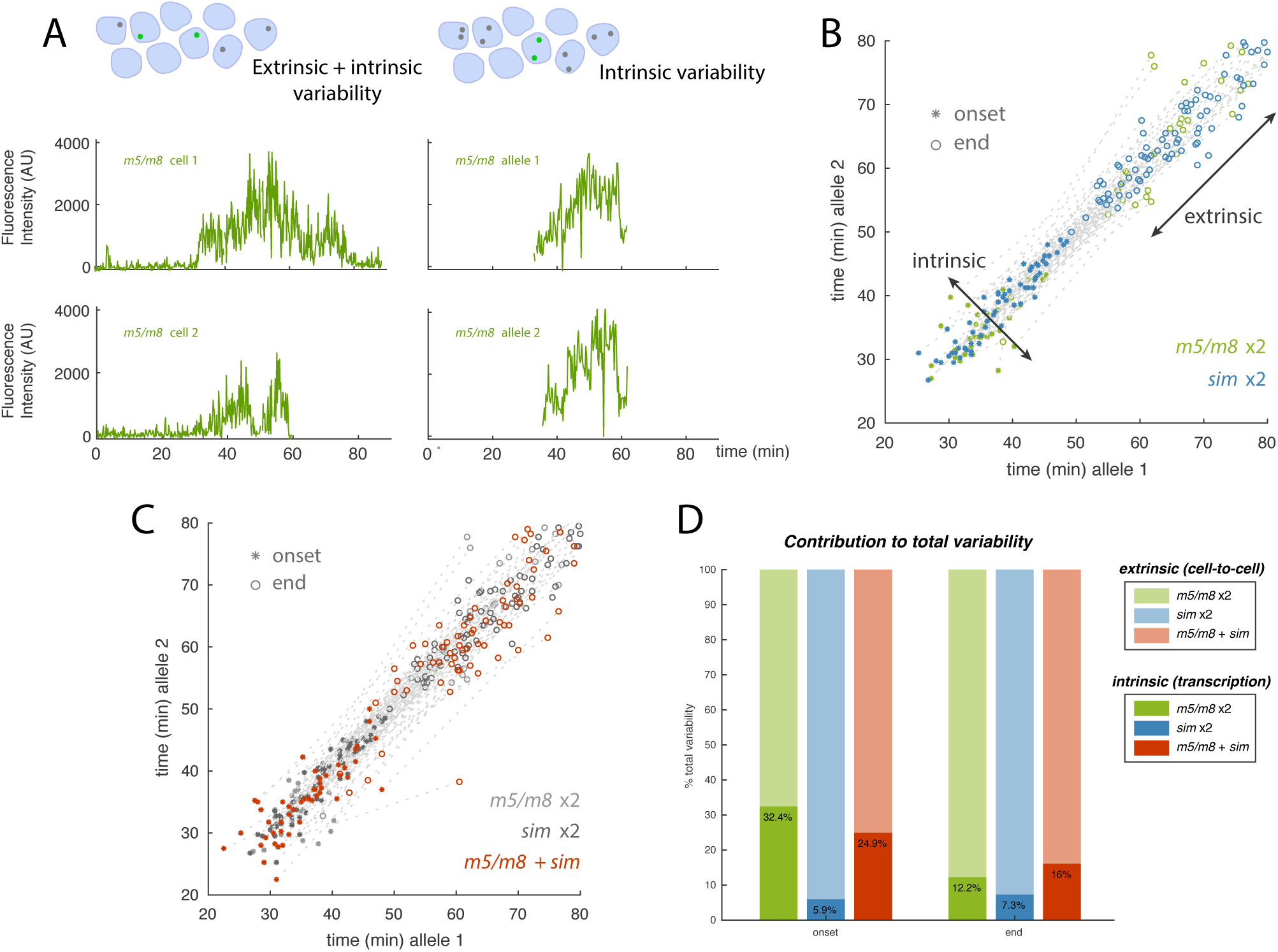
Notch enhancers exhibit low intrinsic variability. **A**) Examples of transcription profiles from *m5/m8* in different nuclei (left panels) and from two alleles in the same nucleus (right panels). **B**) Onset and end-points of activity from individual puncta in nuclei carrying two alleles of *m5/m8* or *sim*. Distribution across the diagonal reflects intrinsic variability (within cells) whereas distribution along the diagonal reflects extrinsic variability (between cells). **C**) Onset and end-points of activity from individual puncta in nuclei carrying an allele of *m5/m8* and an allele of *sim* (data from the individual enhancers, **C**, shown in grey for comparison). **D**) Contribution of intrinsic variability (dark shading) to variability in transcription onset and end-times in the indicated two-allele combinations. Connecting grey lines indicates onset and end times from the same nucleus. The *peve* promoter was used in all reporters unless otherwise specified. n = 2 (*m5/m8* x2), 3 (*sim*x2), 3 (*m5/m8* + *sim*) embryos.

Although the overall temporal profiles of transcription from two alleles in the same cell were similar, the fine grained spikes and troughs were not synchronised (Fig. 2**A**), in agreement with the expectation that transcription from two different loci is largely uncorrelated (Harper et al. 2011; Little et al. 2013; Fritzsch et al. 2018). However, the fluorescent intensities of two alleles at any time point displayed a small but significant positive correlation (R^2^ ~ 0.35), compared to a null correlation when these pairs are randomly assigned (Fig. S2**B**). This argues that the enhancers at the two alleles operate independently while being co-ordinated by the same extrinsic information. Even when *m5/m8* and *sim* were placed in trans in the same cell, there was comparatively little variation in onset times, compared to the variation in different cells (Fig. 2**CD** S2**A**). These results indicate that the properties of *m5/m8* and *sim* ensure they reliably detect extrinsic information in the form of Notch activity, which is initiated within a 5-10 minute time-window, so that their activation is remarkably synchronized within a given nucleus.

### Enhancers detect signal thresholds and signal context

The *m5/m8* and *sim* enhancers appear to act as “persistence detectors”, driving transcription as long as Notch signal(s) are present. They may therefore be simple switches detecting when a signal crosses a threshold (digital encoding) or they could respond in a dose-sensitive manner to Notch activity levels (analog encoding). To distinguish these possibilities, we tested the consequences from additional Notch activity, by supplying ectopic NICD using the *stripe 2* regulatory enhancer from *even-skipped* (*eve2-NICD*). This confers a tightly regulated ectopic stripe of NICD which is orthogonal to the MSE (Fig. 3**A**) (Kosman and Small 1997; Cowden and Levine 2002) and was sufficient to produce ectopic expression from both *m5/m8* and *sim* (**Movie S3**, **Movie S4**).

**Figure 3.**
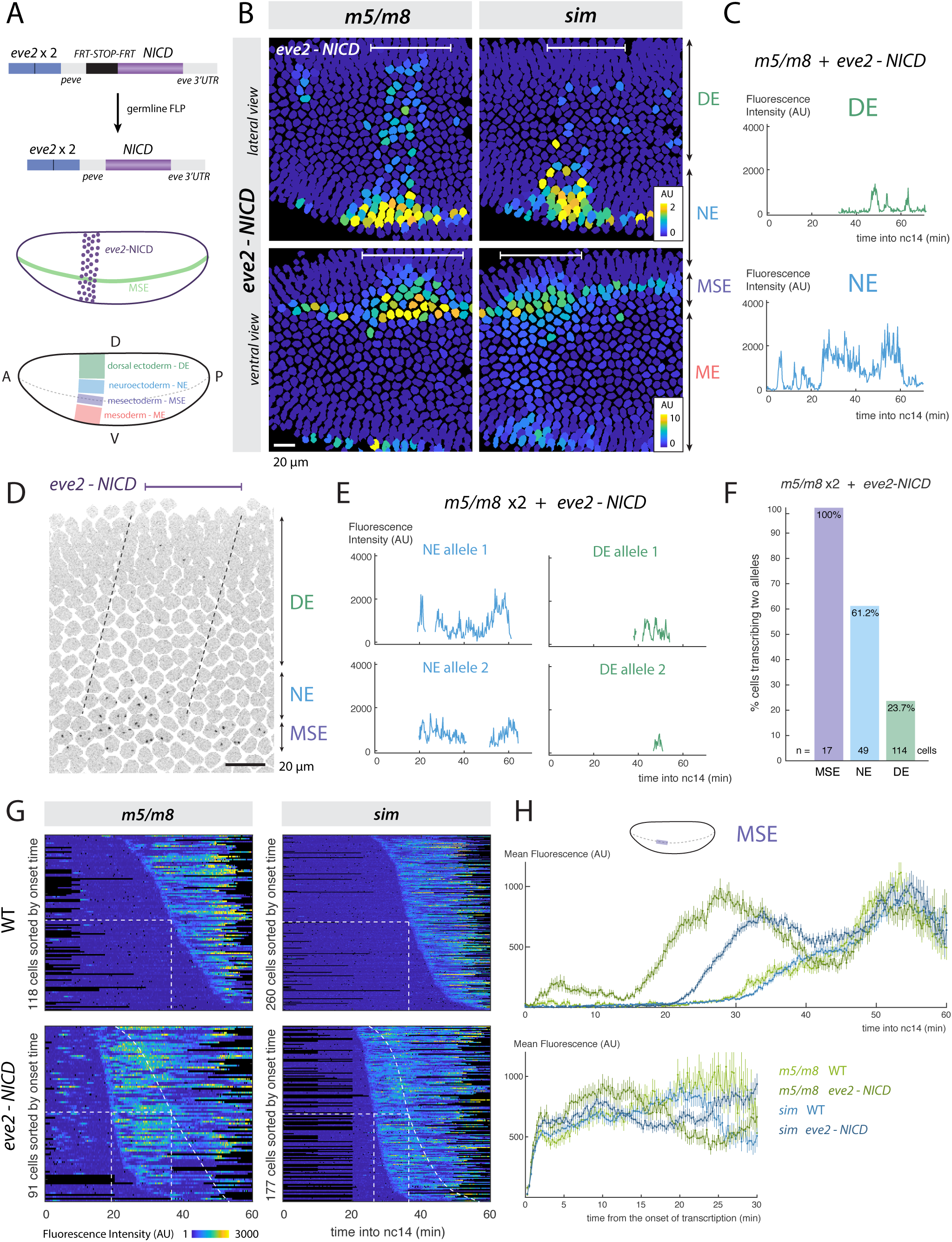
Effects of ectopic NICD on temporal transcription profiles reveals enhancers have different thresholds. **A**) Diagram illustrating the strategy for producing ectopic NICD in a stripe orthogonal to the MSE using *eve2*, with schematics showing expression (purple shading) relative to MSE (green) and DV regions where effects on transcription were quantified. **B**) Still frames of tracked nuclei false-colored for total accumulated signal (note different scales). DE, NE, MSE, ME correspond to the regions shown in **A**. **C**) Illustrative traces from DE (top) and NE (bottom) nuclei, where NICD produces different *m5/m8* transcription profiles. **D**) Still frame of an *eve2-NICD* embryo with two alleles of *m5/m8*. Inverted maximum intensity projection of MCP-GFP channel is overlaid with outlines of tracked nuclei with region of ectopic NICD indicated by dashed lines. **E**) Examples of transcription traces from two *m5/m8* alleles in NE or DE nuclei. **F**) Proportion of nuclei that ever transcribe two *m5/m8* alleles at the same time. **G**) Heatmaps of transcription traces from *m5/m8* and *sim* in MSE nuclei from wild type and *eve2-NICD* embryos, sorted by onset time. Dashed lines indicate onset times in wild type embryos. **H**) Mean profiles of activity in MSE nuclei over time (top) and aligned by onset time (bottom). Aligning by onset time shows that transcription in each nucleus increases steeply in all conditions, indicating that the gradual mean increase reflects small differences in onset times between nuclei. **H** shows mean and SEM of all MSE cells. n = 4 (*m5/m8* WT), 7 (*sim* WT), 6 (*m5/m8 eve2-NICD*), 8 (*sim eve2-NICD*), 4 (*m5/m8* x2 *eve2-NICD*) embryos.

Whereas expression from *m5/m8* and *sim* was almost identical in wild-type embryos, clear differences were revealed by ectopic NICD. First, transcription from *m5/m8* was detected throughout much of the dorsal eve2 domain whereas ectopic transcription from *sim* was only seen in 3-4 cell wide region close to the MSE (Fig. 3**B**), consistent with previous observations (Cowden and Levine 2002; Zinzen et al. 2006a). Second, although both enhancers initiated transcription prematurely, because the ectopic NICD was produced from early nc14 (Bothma et al. 2014), the onset of transcription from *m5/m8* was significantly earlier than that from *sim* (Fig. 3**GH**). Given that both enhancers are exposed to the same temporal pattern of NICD, this difference in their initiation times implies that the two enhancers have different thresholds of response to NICD, with *m5/m8* responding to lower doses and hence being switched-on earlier. Therefore, we hypothesize that *m5/m8* and *sim* respond at the same time in wild-type embryos because the normal ligand-induced signaling leads to a sharp increase in NICD.

We also detected differences in the dynamics of *m5/m8* according to the location of the NICD-expressing nucleus along the DV axis. Nuclei closer to the MSE stripe (in the neuroectoderm, NE) exhibited strong activity, with a temporal pattern that resembled that in the MSE (Fig. 3**C**, bottom). In contrast nuclei in dorsal regions (dorsal ectoderm, DE) underwent resolved bursts of transcriptional activity (Fig. 3**C**, top). Ectopic NICD also induced ‘bursty’ expression from *sim* in the mesoderm (ME), but was not capable of turning on *m5/m8* in that region (**Movie S5**). ‘Bursty’ transcription in the DE was also associated with more stochastic activation. In embryos with two *m5/m8* alleles, both were activated in response to *eve2-NICD* in most MSE and NE nuclei, whereas only a single allele was active at any one time in most DE nuclei (Fig. 3**DF**, **Movie S6**). Furthermore, in the few DE nuclei where both alleles became active, there was greater variability in onset times and the profiles were less coordinated (Fig. 3**E**, S2**D**). The positional differences in dynamics suggest that intrinsic cellular conditions, likely the expression levels of specific transcription factors, influence the way that enhancers “read” the presence of NICD. Such factors must therefore have the capability to modulate the dynamics of transcription.

### Notch activity tunes transcription burst size

To further test how Notch responsive enhancers respond to doses of signal, we introduced a second *eve2-NICD* transgene. MSE transcription from *sim* in the presence of 2x*eve2-NICD* initiated earlier and achieved higher levels than with 1x*eve2-NICD* (Fig. 4**A**, left). This is consistent with the hypothesis that *sim* responds to thresholds of NICD concentration, as the cells will reach a given concentration of signal more quickly in embryos with 2x*eve2-NICD*. The mean levels of transcription increased in ME as well as in MSE regions (Fig. 4**A-C**), further indicating a dose-sensitive response. In contrast, the levels and onset of MSE transcription from *m5/m8* did not significantly change in 2x*eve2-NICD* embryos (Fig. 4**A**, right). This saturation in output levels of transcription from *m5/m8* only occurred in the MSE, as the more stochastic activity in the DE remained sensitive to increases in NICD, being detected in a greater proportion of cells and over longer periods (Fig. S4**A**).

**Figure 4.**
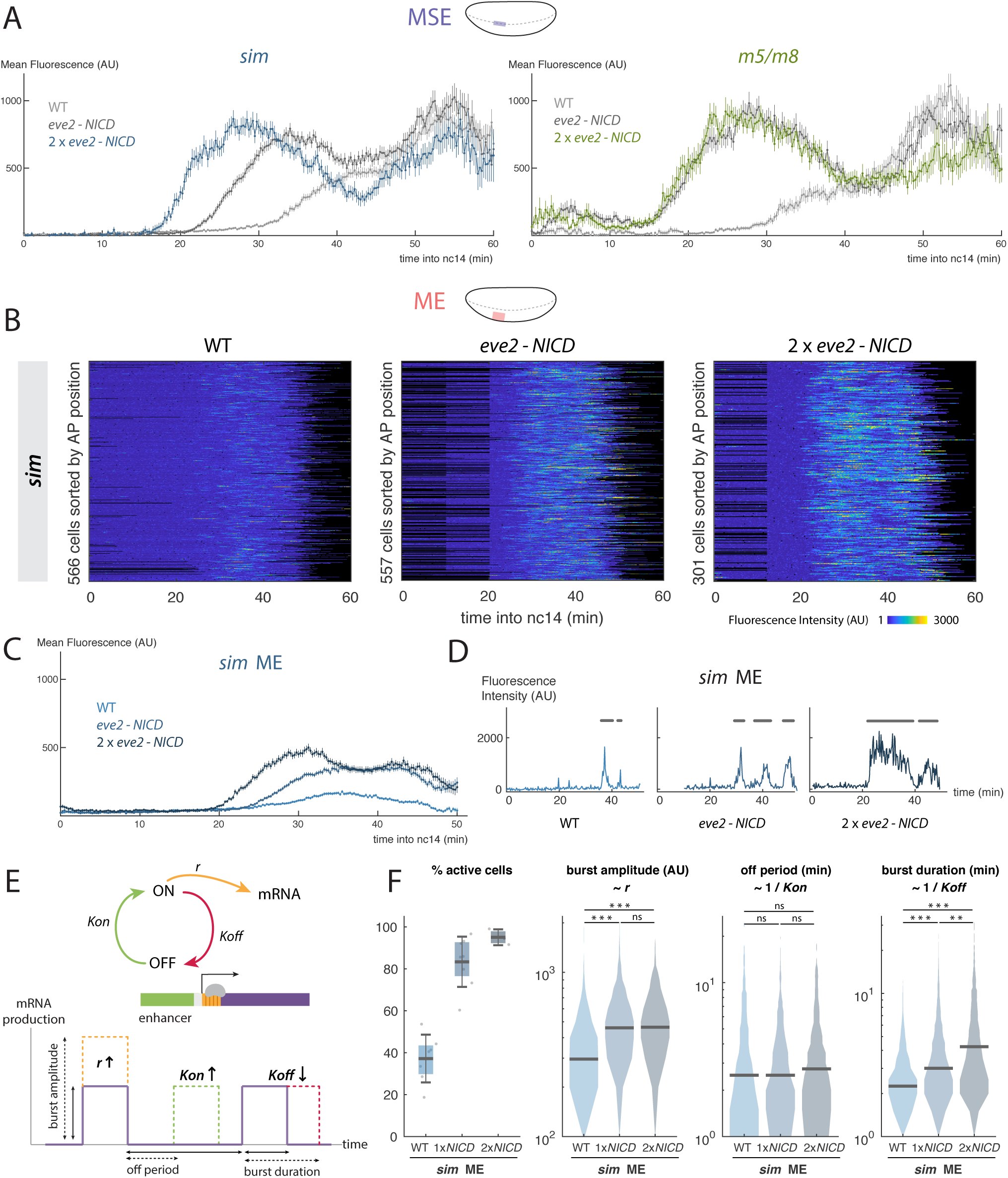
Notch produces a dose-sensitive response by regulating transcription burst size. **A**) Mean levels of transcription from *sim* (left) and *m5/m8* (right) in the MSE with an additional NICD insertion (2x*eve2-NICD*), compared to 1x*eve2-NICD* and wild type. **B**) Heatmaps depicting *sim* activity in ME nuclei in three conditions as indicated. Note the different scale range compared to Fig. 3**G**. **C**) Increased mean levels of transcription from *sim* in the ME produced by different doses of NICD. **D**) Examples of transcription traces from single ME nuclei in wild type, 1x*eve2-NICD* and 2x*eve2-NICD*. Burst periods are marked with a grey line. **E**) Schematic of the model: an enhancer cycles between ON and OFF states and produces mRNA when ON. Changes in bursting amplitude, off period and bursting duration correlate with changes in kinetic constants *r*, *Kon* and *Koff*. **F**) Quantification of individual burst properties from *sim* in ME of wild type, 1x*eve2-NICD* and 2x*eve2-NICD* embryos. Boxplots indicate median, with 25-75 quartiles; error bars are SD. Violin plots, distributions of the analyzed bursts, bar indicates the median. In **A** and **C** mean fluorescence values and SEM are plotted. Grey lines are reproduced from Fig. 3**H**. n cells for **B-F** are indicated in **B**. Differential distributions tested with two-sample Kolmogorov-Smirnov test: pvalues <0.01(*), <10^−5^(**), <10^−10^(***). n = 3 (*m5/m8* 2x*eve2-NICD*), 3 (*sim* 2x*eve2-NICD*) embryos.

To distinguish different models for how NICD confers a dose-sensitive response, we took two strategies to analyze its effect on transcriptional bursting dynamics and focused on regions where individual bursts of transcription were resolved. Both approaches assume a two state model where the enhancer is switched between an OFF and ON state with switching rates *Kon* and *Koff* and confers transcription initiation rate *r* in the ON state (Fig. 4**E**)(Peccoud and Ycart 1995; Larson et al. 2009). In the first approach we directly measured bursting amplitude, off period between bursts and bursting length as approximations for *r*, *Kon* and *Koff*, respectively (Fig. 4**E**). In most previous enhancers analyzed in this way, the off period is the most affected, leading to changes in the bursting frequency (Fukaya et al. 2016; Fritzsch et al. 2018; Lammers et al. 2018). However, when we quantified the effect from different doses of NICD on *sim* in the ME we found that the bursting length consistently increased with higher amounts of NICD whereas the off period between bursts remained constant (Fig. 4**DF**). This indicates that the main effect of NICD is to keep the enhancer in the ON state for longer - ie. decreasing *Koff* - rather than increasing the frequency with which it becomes active - i.e. increasing *Kon*. The bursting amplitude also increased with 1x*eve2-NICD* but this was not further enhanced by 2x*eve2-NICD* (Fig. 4**DF**). Overall therefore, increasing levels of NICD in the ME result in *sim* producing an increase in transcription burst size (duration x amplitude) rather than an increase in the frequency of bursts. Transcription in other regions and enhancers (*m5/m8* DE and *m8NE* ME) showed similar increase in burst size in response to the dose of NICD (Fig. S4**A-C**) suggesting this is a general property of these Notch responsive enhancers.

We developed a second approach, based on the noise properties of transcription, to analyze the changes in the dynamics even where single bursts of activity could not be defined. To do so, we used a mathematical model of transcription to account for the initiating mRNA molecules (Fig. S3**A**). Using derivations from the mathematical model and testing them in simulations, we looked for signatures that would be produced if the mean of initiating mRNAs (equivalent to the mean fluorescence from MS2 puncta) were increasing due to changes in *r*, *Kon* or *Koff*. This showed that the effects on the Fano factor ratio between the two conditions and on their autocorrelation function (ACF) could be used to correctly predict which of the three parameters could account for an increase in the mean (Fig. S3**B**, Supplementary Methods). First we tested the modelling approach with the data from the promoter swap experiments. Analyzing the differences in the mean indicated that they are most likely due to increases in *r* (Fig. S4**D**), as expected if promoters influence the rate of polymerase release but not enhancer activation per se. When we then applied the model to data from the transcription profiles produced by different doses of NICD in the ME, results were most compatible with the causal effect being an increase in *r* or a decrease in *Koff* (Fig. S4**E**) depending on which two conditions were compared. Thus this second approach also indicated that NICD elicits an increase in burst size rather than in burst frequency. Both approaches therefore converge on the model that, above the critical threshold level of NICD, further increases in NICD levels prolong the period that each enhancer remains in the ON state.

Finally, we used an enhancer - promoter combination that produced higher mean levels (*m5/m8-pm5*, Fig. S1**E**) to investigate whether the saturation that occurred with ectopic NICD was due to the *peve* promoter having achieved maximal initiation rate. Strikingly, the substitution of *pm5* for *peve* did not result in significantly higher maximal levels in the presence of *eve2-NICD* (Fig. S4**F**) although it did in wild-type embryos (Fig. S1**E**). This indicates that the saturation of the response that occurs with higher levels of NICD stems from the *m5/m8* enhancer rather than the promoter and argues that enhancers reach a maximal “ON” state that they cannot exceed even if more NICD is provided.

### Paired CSL motifs augment burst-size not threshold detection

The *m5/m8* and *sim* enhancers both respond to NICD but they initiate transcription at different thresholds. How is this encoded in their DNA sequence? A prominent difference between the two is that *m5/m8* contains a paired CSL motif (so-called SPS motif) whereas *sim* does not (Fig. S5**A**). To test their role, we replaced two CSL motifs in *sim* with the SPS motif from *m5/m8* and conversely perturbed the SPS in *m5/m8* by increasing the spacing between the two CSL motifs (Fig. S5**A**). As SPS motifs permit co-operative binding between two NICD complexes, we expected that enhancers containing an SPS motif (*sim^SPS^* and *m5/m8*) would exhibit earlier onsets of activity than their cognates without (*sim* and *m5/m8^insSPS^*). However this was not the case for either *sim* and *sim^SPS^* (Fig. 5**AB**) or *m5/m8* and *m5/m8^insSPS^* in either wild type or *eve2-NICD* embryos (Fig. S5**DE**). These profiles suggest that the SPS motifs are not responsible for the difference in the threshold levels of NICD required for *m5/m8* and *sim* activation.

**Figure 5.**
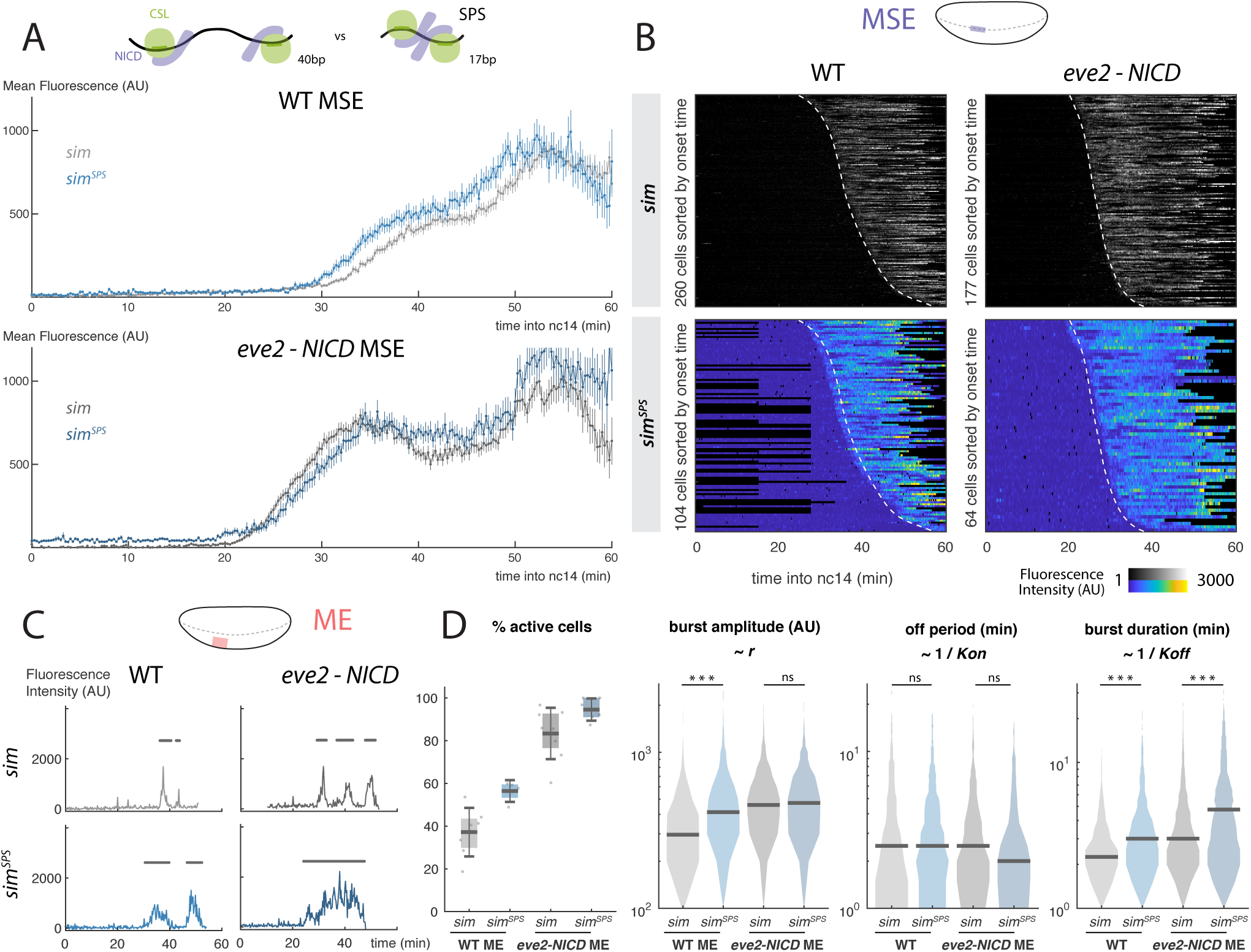
Optimized Su(H) motif organization enhances bursting size. **A**) Mean levels of transcription in MSE nuclei when two Su(H) motifs in *sim* are replaced with an optimal paired SPS motif (*sim^SPS^*) in wild type (top) and *eve2-NICD* (bottom) embryos. Mean and SEM shown. **B**) Heatmaps of transcription in MSE nuclei from *sim^SPS^* and *sim* in wild-type and 1x*eve2-NICD* embryos, sorted by onset time. Dashed lines indicate onset times for un-mutated *sim*. **C**) Examples of fluorescent traces from *sim* and *sim^SPS^* in ME nuclei. Burst periods are indicated with grey lines. **D**) *sim^SPS^* is active in a higher proportion of cells and with increased burst size compared to *sim*. Boxplots indicate median, 25-75 quartiles and errorbars are SD. Violin plots, distribution for all bursts measured in the ME, bar indicates the median. Differential distributions tested with two-sample Kolmogorov-Smirnov test: pvalues <0.01(*), <10^−5^(**), <10^−10^(***). n = 4 (*sim^SPS^* WT) and 6 (*sim^SPS^ eve2-NICD*) embryos. Grey lines, heatmaps and violin plots are re-plotted from Fig. 3**GH** and 4**DF** for comparison.

Changes to the CSL motifs did however affect the mean levels of activity. *sim^SPS^* directed higher mean levels of activity compared to *sim* in both wild type and *eve-NICD* embryos (Fig. 5**A** S5**B**). Conversely, *m5/m8^insSPS^*directed lower levels compared to *m5/m8* (Fig. S5**D**). Analysing the traces from *sim* in the ME, where cells undergo resolved bursts of transcription, revealed that the SPS motif (*sim^SPS^*) led to larger burst-sizes - i.e. increased the amplitude and the duration - compared to the wild type *sim* enhancer (Fig. 5**CD**). Conversely, the continuous profile produced by *m5/m8* in the MSE was broken into smaller bursts when the SPS was disrupted (Fig. S5**FG**). The effects on bursting size are similar to those seen when the dose of NICD was altered, suggesting that enhancers containing SPS sites respond to a given level of NICD more effectively. They do not however appear to affect the amount of NICD required for their initial activation, i.e. the threshold required for the enhancer to be switched on. This implies that burst-size modulation and response threshold can be uncoupled and potentially could be encoded independently at the DNA level.

### Regional factors prime enhancers for fast and sustained activation

Under ectopic NICD conditions, *m5/m8* and *sim* both produce sustained transcription profiles in the MSE and NE, whereas elsewhere they generate stochastic and “bursty” transcription. This suggests that other factors are “priming” the enhancers to respond to NICD. Good candidates are the factors involved in DV patterning at this stage, the bHLH transcription factor Twist (Twi) and/or the Rel protein Dorsal (dl). Indeed, the region where the enhancers generate sustained profiles in response to *eve2-NICD* overlaps the domain of endogenous Twist and Dorsal gradients (Fig S6**B**)(Zinzen et al. 2006b). Furthermore, *m5/m8* and *sim* both contain Twist and Dorsal binding motifs (Fig. S6**A**) and previous studies indicated that Twist is important for *sim* activity although it was not thought to contribute to *m5/m8* activity (Zinzen et al. 2006a).

To test if Twist and Dorsal are responsible for the different dynamics of *m5/m8* transcription observed in the MSE and DE with ectopic NICD (Fig. 3**C**), we mutated the Twist and/or Dorsal binding motifs in *m5/m8* (Fig. S6**A**). Strikingly, mutating the three Twist or two Dorsal motifs produced a delay in the start of transcription in both wild-type and *eve2-NICD* embryos. These effects were even more pronounced when both Twist and Dorsal motifs were mutated together (Fig. 6**B**), implying that, without Twist or Dorsal, *m5/m8* requires a higher threshold of NICD to become active or responds more slowly to the same threshold. The mean transcription levels were also reduced in all cases (Fig. 6**A**).

**Figure 6.**
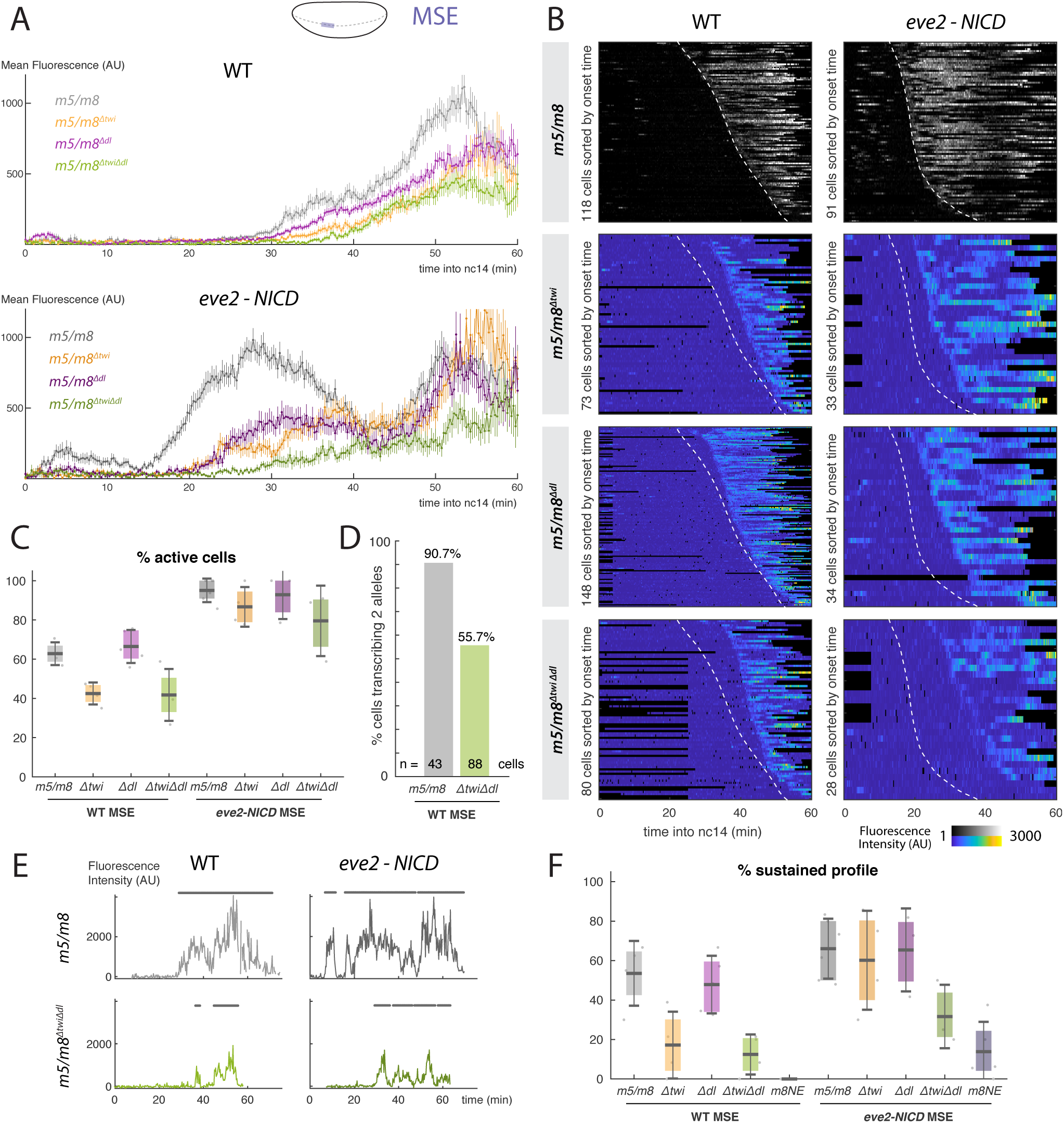
Twist and Dorsal prime the response of *m5/m8* to NICD. **A**) Mean levels of activity in wild type (top) and *eve2-NICD* (bottom) produced by mutations in Twist and/or Dorsal binding motifs in *m5/m8*. **B**) Heatmaps of mutated enhancer activity in MSE nuclei in wild type and *eve2-NICD*, sorted by onset time. Dashed lines indicate onset times for un-mutated enhancer. **C**) Proportion of active cells in the MSE in wild type and *eve2-NICD* embryos when Twist and/or Dorsal motifs are mutated, compared to unmutated *m5/m8*. **D**) Proportion of active cells transcribing two alleles at any point, in embryos containing two copies of *m5/m8* or of *m5/m8*^Δ*twi*Δ*dl*^. **E**) Examples of transcription traces from wild type and mutated *m5/m8* in MSE nuclei from wild type and *eve2-NICD* embryos. Profiles from *m5/m8*^Δ*twi*Δ*dl*^ MSE cells exhibit ‘bursty’ transcription. ON periods are marked with a grey line. **F**) Proportion of MSE cells per embryo displaying a sustained profile of transcription, defined by one burst of >10 min. Median, quartiles and SD are shown. Grey lines and heatmaps are re-plotted from Fig. 3**GH**. n = 4 (*m5/m8*^Δ*twi*^ WT), 5 (*m5/m8*^Δ*dl*^ WT), 4 (*m5/m8*^Δ*twi*Δ*dl*^ WT), 4 (*m5/m8*^Δ*twi*^ *eve2-NICD*), 3 (*m5/m8*^Δ*dl*^ *eve2-NICD*), 3 (*m5/m8*^Δ*twi*Δ*dl*^ *eve2-NICD*), 3 (*m8NE* WT), 5 (*m8NE eve2-NICD*), 3 (*m5/m8*^Δ*twi*Δ*dl*^ x2).

Mutating the Twist motifs had two additional effects: the overall proportion of active cells in the MSE was reduced (Fig. 6**C**) and out of those active, fewer exhibited the sustained profile observed with wild type *m5/m8* (Fig. 6**EF**). Instead most cells displayed a ‘bursty’ transcription profile (Fig. 6**E**), similar to those elicited by NICD in the DE. Although the mutated Twist motifs led to bursty profiles in wild type embryos, these effects were partially rescued when ectopic NICD was provided (Fig. 6**CF**, S6**C**). However, when both Dorsal and Twist motifs were mutated, the proportions of active nuclei and of nuclei with sustained profiles decreased even in the presence of ectopic NICD (although mutation of Dorsal motifs alone did not produce a significant decrease in either property) (Fig. 6**CF**, S6**C**). The decrease in the overall proportion of active cells suggests that Twist and Dorsal regulate the probability of *m5/m8* to activate transcription in response to Notch. In agreement, in embryos with two alleles of a reporter, the proportion of cells transcribing from both alleles was much lower for *m5/m8*^Δ*twi*Δ*dl*^ than for *m5/m8* (Fig. 6**D**). Additionally, in those nuclei where both reporters were active, there was considerably more variability in the onset times for *m5/m8*^Δ*twi*Δ*dl*^ compared to *m5/m8* (Fig. S6**E**). The results are therefore consistent with a role for Twist and Dorsal in priming the *m5/m8* enhancer to become active in response to Notch and produce sustained activity. In their absence the ability of *m5/m8* to initiate transcription becomes much more stochastic. Likewise, *m8NE*, (which contains only a single Twist motif, Fig. S6**A**) also produced delayed and ‘bursty’ rather than sustained transcription (Fig. 6**F**, S6**D**). The two MSE enhancers thus appear to be especially configured to respond in a fast and sustained manner at this stage.

## Discussion

Developmental signaling pathways have widespread roles but currently we know relatively little about how the signaling information is decoded to generate the right transcriptional outcomes. We set out to investigate principles that govern how Notch activity is read by target enhancers in the living animal, using the MS2/MCP system to visualize nascent transcripts in *Drosophila* embryos and focusing on two enhancers that respond to Notch activity in the MSE. Three striking characteristics emerge. First, MSE enhancers are sensitive to changes in the levels of NICD, which modulate the transcriptional burst size rather than increasing burst frequency. Second, the activation of both MSE enhancers is highly synchronous. Indeed, within one nucleus the two enhancers become activated within few minutes of one another. Third, both MSE enhancers confer a sustained response in the wild-type context. This synchronized and persistent activity of the MSE enhancers contrasts with the stochastic and bursty profiles that are characteristics of most other enhancers that have been analyzed (Little et al. 2013; Fukaya et al. 2016; Fritzsch et al. 2018) and relies on the MSE enhancers being “primed” by regional transcription factors Twist and Dorsal. We propose that such priming mechanisms are likely to be of general importance for rendering enhancers sensitive to signals so that a rapid and robust transcriptional response is generated.

### Priming of enhancers sensitizes the response to NICD

Transcription of most genes in animal cells occurs in bursts interspersed with refractory periods of varying lengths, that are thought to reflect the kinetic interactions of the enhancer and promoter (Bartman et al. 2016). However, the MSE enhancers appear to sustain transcription for 40-60 minutes, without detectable periods of inactivity, albeit that very short off-periods might not have been resolved by our assays. Calculation of the autocorrelation function in traces from these nuclei suggest very slow transcriptional dynamics (Fig. S4**ED**) (Desponds et al. 2016; Lammers et al. 2018), which would be consistent with one long period of activity as opposed to overlapping short bursts. This fits with a model where promoters can exist in a permissive active state, during which many “convoys” of polymerase can be fired without the promoter reverting to a fully inactive condition (Tantale et al. 2016). The rapid successions of initiation events are thought to require Mediator complex (Tantale et al. 2016), which was also found to play a role in the NICD-mediated increase in residence time of CSL complexes (Gomez-Lamarca et al. 2018). We propose that sustained transcription from *m5/m8* and *sim* reflects a switch into a promoter permissive state, in which general transcription factors like Mediator remain associated with the promoter so long as sufficient NICD is present, allowing repeated re-initiation.

However, the ability to drive fast and sustained activation is not a property of NICD itself. For example, when ectopic NICD was supplied, cells in many regions of the embryo responded asynchronously and underwent short bursts of activity. Furthermore, variable and less sustained cell-by-cell profiles were generated in the MSE region when the binding motifs for Twist and Dorsal in *m5/m8* were mutated. The presence of these regional factors appears to sensitize the enhancers to NICD, a process we refer to as enhancer priming. This has two consequences. First, it enables all nuclei to respond rapidly to initiate transcription in a highly coordinated manner once NICD reaches a threshold level. Second, it creates an effective ‘state transition’ so that the presence of NICD can switch the promoter into a permissive condition to produce sustained activity (Fig. 7). We propose a priming mechanism, rather than classic co-operativity, because Twist and Dorsal alone are insufficient to drive any enhancer activity. Furthermore, since the enhancers immediately achieve sustained activity when NICD is produced, it is most likely that Twist and Dorsal are required prior to the recruitment of NICD, although both may continue to play a role independently of priming after transcription is initiated, as suggested by the lower mean levels obtained when only Twist or Dorsal motifs are mutated. Another contributory factor may be recruitment of the co-repressor complex containing CSL and Hairless, whose presence at primed enhancers could poise for activation and set the threshold (Barolo and Posakony 2002).

**Figure 7.**
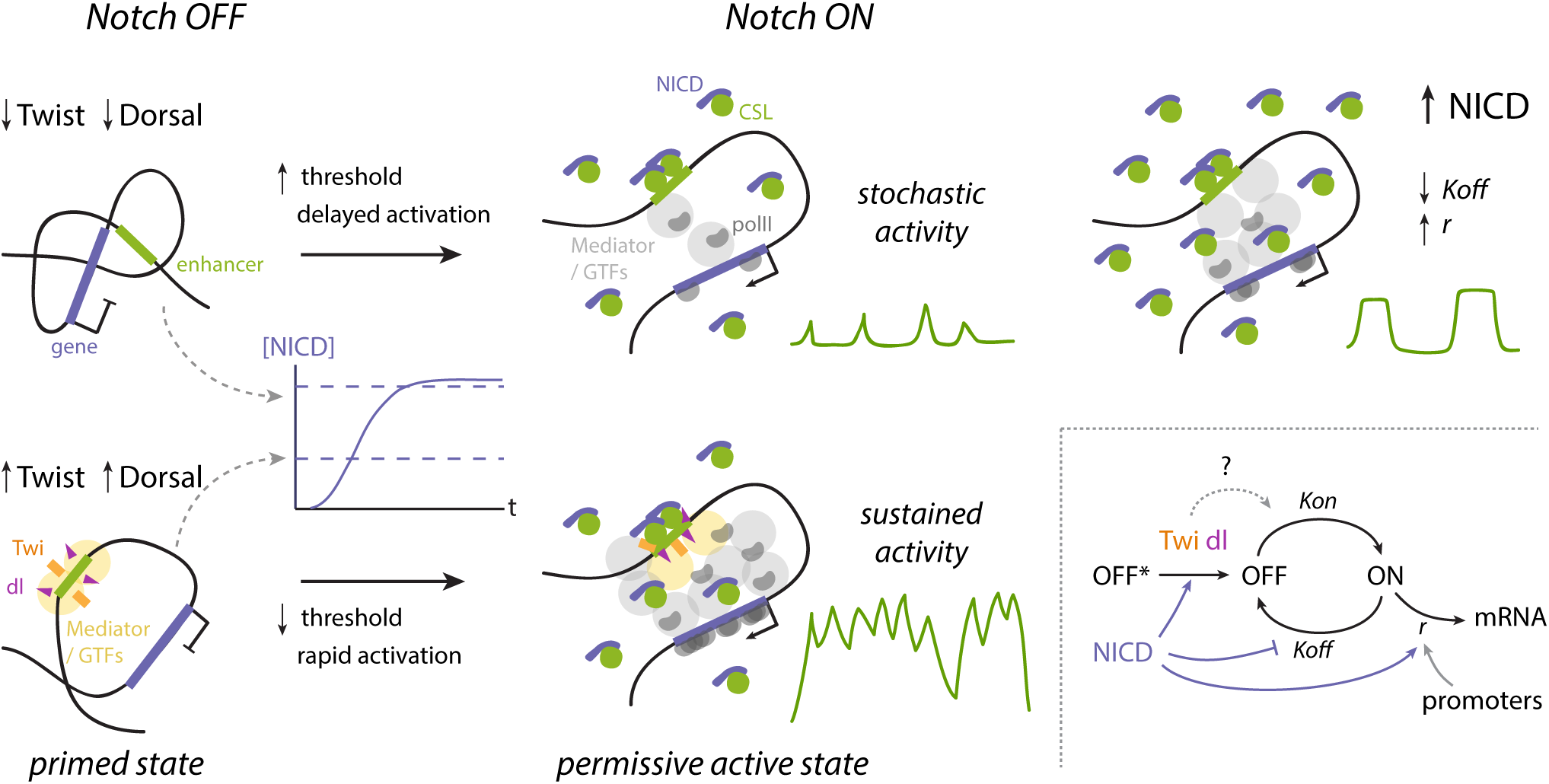
Model of transcriptional regulation by Notch through enhancer priming and burst size modulation. Priming by the tissue-specific factors Twist and Dorsal produces rapid activation in response to NICD and a state transition into a permissive active state in which sustained transcription can be produced without cycling between ON and OFF states. In the absence of these factors stochastic activity is produced in response to NICD. Increasing levels of NICD regulate the overall probability of the enhancer switching on (OFF* to OFF, which is also modulated by Twist and Dorsal), and increase the bursting size (higher *r* and lower *K_off_*). In contrast different promoters control the initiation rate *r* but do not affect enhancer activation dynamics. The effects of Twist and Dorsal on enhancer priming might also act by modulating the same parameters of transcription.

Our explanation that the synchronous activation of the MSE enhancers reflects their requirements for a critical concentration of NICD is borne out by their responses when levels of NICD are increased. Notably, while *sim* and *m5/m8* had almost identical dynamics in wild-type embryos, their response to ectopic NICD differed, suggesting that they detect different thresholds. Indeed, doubling the dose of ectopic NICD further accelerated onset times of *sim* in agreement with the model that the enhancers detect NICD levels. Threshold detection does not appear to rely on the arrangement of CSL motifs, as onset times of *m5/m8* or *sim* were unaffected by changes in the spacing of CSL paired sites. In contrast, mutating Twist or Dorsal binding-motifs in *m5/m8* delayed the onset, arguing that these factors normally sensitize the enhancer to NICD, enabling responses at lower thresholds.

We propose that enhancer priming will be widely deployed in contexts where a rapid and consistent transcriptional response to signaling is important, as in the MSE where a stripe of cells with a specific identity is established in a short time-window. In other processes where responses to Notch are more stochastic, as during lateral inhibition, individual enhancers could be preset to confer different transcription dynamics. This appears to be the case for a second enhancer from *E(spl)-C* (*m8NE*) which generates a stochastic response in the MSE cells, similar to that seen for the MSE enhancers when Twist and Dorsal sites are mutated. This illustrates that the presence or absence of other factors can toggle an enhancer between conferring a stochastic or deterministic response to signalling.

### NICD regulates transcription burst size

Manipulating NICD levels revealed that Notch responsive enhancers act as analog devices that can measure and broadcast variations in levels. Increased NICD levels have a consistent effect on enhancer activity irrespective of the priming state of the enhancer: an increase in burst size. This can be most readily quantified in regions where NICD elicits discrete bursts of transcription initiation, such as the DE for *m5/m8* or ME for *sim* and *m8NE*. Transcriptional bursting has been formalized as a two-state model where the promoter toggles between ON and OFF states, conferring a transcription initiation rate when ON (Peccoud and Ycart 1995; Larson et al. 2009). Changes in the duration or frequency of the bursts lead to an overall increase in transcription. Most commonly, differences in enhancer activity have been attributed to changes in the switching on probability (*Kon*), leading to changes in burst frequency (Larson et al. 2013; Senecal et al. 2014; Fukaya et al. 2016; Fritzsch et al. 2018; Lammers et al. 2018; Berrocal et al. 2018). We were therefore surprised to find that higher doses of NICD did not increase burst frequency. Instead they produced bigger bursts, both by increasing bursting amplitude, equivalent to the rate of transcription initiation, and bursting length, indicative of the total time the enhancer stays in the ON state. Modifications to the CSL motifs also impacted on the same parameters. Thus, enhancers with paired motifs (SPS), which favour NICD dimerization (Nam et al. 2007), produced larger transcription bursts than those where the motifs are further apart. This suggests that paired motifs can ‘use’ the NICD present more efficiently, potentially because optimally configured sites increase the likelihood that at least one NICD will be bound at any time. Interestingly, even though *m5/m8* and *sim* contain different arrangements and numbers of CSL motifs they have converged to produce the same mean levels of transcription in wild type embryos.

Two models would be compatible with the observations that effective NICD levels alter the burst size. In the first model, increasing the concentration of NICD when the enhancer is activated would create larger PolII clusters. This is based on the observation that low complexity activation domains in transcription factors can form local regions of high concentration of transcription factors, so-called “hubs”, which in turn are able to recruit PolII (Mir et al. 2017; Tsai et al. 2017; Lu et al. 2018). As the lifetime of PolII clusters appears to correlate with transcriptional output (Cho et al. 2016), the formation of larger PolII clusters would in turn drive larger bursts. In the second model, NICD would be required to keep the enhancer in the ON state, for example by nucleating recruitment of Mediator and/or stabilizing a loop between enhancer and promoter, which would in turn recruit PolII in a more stochastic manner. General factors such as Mediator have been shown to coalesce into phase-separated condensates that compartmentalize the transcription apparatus (Cho et al. 2018; Sabari et al. 2018; Boija et al. 2018) and these could form in a NICD dependent manner. Whichever the mechanism, persistence of the clusters/ON state requires NICD yet must be compatible with NICD having a short-lived interaction with its target enhancers (Gomez-Lamarca et al. 2018). Furthermore, the fact that the activity of *m5/m8* saturates with one *eve2-NICD* construct, and can’t be enhanced by providing a more active promoter, suggests that there is a limit to the size or valency of the clusters that can form.

Although unexpected, the ability to increase burst size appears to be a conserved property of NICD. Live imaging of transcription in response to the Notch homologue, GLP-1, in the *C.elegans* gonad also shows a change in burst size depending on the signalling levels (Lee et al. 2018). As the capability to modulate burst size is likely to rely on the additional factors recruited, the similarities between the effects in fly and worm argue that a common set of core players will be deployed by NICD to bring about the concentration-dependent bursting properties.

## Author contributions

JFS and SJB planned the experiments; JFS conducted the experiments; JFS,NL,HG developed the computational modelling and analysis; JFS, SJB wrote the manuscript; NL,HG edited the manuscript.

## Acknowledgments

We thank members of the Bray Lab and of the Notch community for helpful discussions and Bill Harris and Maria J. Gomez-Lamarca for comments on the manuscript. Thanks to the Sanson, Small and St Johnston, labs for providing flies and plasmids and to Kat Millen and the Genetics Fly Facility for injections. This work was supported by a Programme grant from the Medical Research Council to SJB and by a PhD studentship to JFS from the Wellcome Trust (109144/Z/15/Z). HGG was supported by the Burroughs Wellcome Fund Career Award at the Scientific Interface, the Sloan Research Foundation, the Human Frontiers Science Program, the Searle Scholars Program, the Shurl & Kay Curci Foundation, the Hellman Foundation, the NIH Director’s New Innovator Award (DP2 OD024541-01), and an NSF CAREER Award (1652236). NL was supported by NIH Genomics and Computational Biology training grant 5T32HG000047-18. We also want to thank the Physical Biology of the Cell Course at the Marine Biological Laboratory (Woods Hole, MA), where the modelling approached used in this work developed.

## STAR Methods

### CONTACT FOR REAGENT AND RESOURCE SHARING

Further information and requests for resources and reagents should be directed to and will be fulfilled by the Lead Contact, Sarah J. Bray.

### EXPERIMENTAL MODEL AND SUBJECT DETAILS

#### Experimental Animals

*Drosophila melanogaster* flies were grown and maintained on food consisting of the following ingredients: Glucose 76g/l, Cornmeal flour 69g/l, Yeast 15g/l, Agar 4.5g/l, Methylparaben 2.5ml/l. Embryos were collected on apple juice agar plates with yeast paste. Animals of both sexes were used for this study.

#### Cloning and transgenesis

##### Generation of MS2 reporter constructs

*MS2* loops were inserted upstream of a *lacZ* transcript within the 5’UTR and then the resulting reporter was combined with different enhancers and promoters. 24 *MS2* loops were cloned from *pCR4-24XMS2SL-stable* (Addgene #31865) into *pLacZ2-attB* (Bischof et al. 2013) using *Eco*RI sites. The *m5/m8*, *sim* and *m8NE* enhancers (Zinzen et al. 2006a; Kramatschek and Campos-Ortega 1994) were amplified from genomic DNA and cloned into *pattB-MS2-LacZ* using *Hin*dIII/*Age*I sites (primers in Table S1). Subsequently the promoters *hsp70*, *peve*, *pm5*, *pm6*, *pm7*, *pm8* and *psimE* were cloned by Gibson Assembly (Gibson 2011) in *pattB-m5/m8-MS2-LacZ*, *pattB-sim-MS2-LacZ* and/or *pattB-m8NE-MS2-LacZ* (primers in Table S1) using the *Age*I restriction site and incorporating a *Eag* I site.

Su(H), Twi, dl and Sna binding motifs were identified using ClusterDraw2 using the PWM from the Jaspar database for each transcription factor. Motifs with scores higher than 6 and pvalues 0.001 were selected. Primers to create *sim^SPS^*, *m5/m8^insSPS^*, *m5/m8*^Δ*twi*^, *m5/m8*^Δ*dl*^ and *m5/m8*^Δ*twi*^ ^Δ*dl*^ are detailed in Table S1. All mutations were first introduced by Gibson Assembly in the enhancers contained in *pCR4* plasmids and then transferred to *pattB-peve-MS2-lacZ* using *Hin*dIII and *Age*I sites.

The following constructs have been generated and inserted by ΦC31 mediated integration (Bischof et al. 2007) into an *attP* landing site in the second chromosome – *attP40*, 25C – to avoid positional effects in the comparisons: *pattB-m5/m8-peve-MS2-LacZ*, *pattB-m5/m8-hsp70-MS2-LacZ*, *pattB-m5/m8-pm5-MS2-LacZ*, *pattB-m5/m8-pm6-MS2-LacZ*, *pattB-m5/m8-pm7-MS2-LacZ*, *pattB-m5/m8-pm8-MS2-LacZ*, *pattB-m5/m8-psimE-MS2-LacZ*, *pattB-m8NE-peve-MS2-LacZ*, *pattB-sim-peve-MS2-LacZ*, *pattB-sim-psimE-MS2-LacZ*, *pattB-sim^SPS^-peve-MS2-LacZ*, *pattB-m5/m8^insSPS^-peve-MS2-LacZ*, *pattB-m5/m8*^Δ*twi*^*-peve-MS2-LacZ*, *pattB-m5/m8*^Δ*dl*^*-peve-MS2-LacZ* and *pattB-m5/m8*^Δ*twi*^ ^Δ*dl*^*-peve-MS2-LacZ*. *pattB-m5/m8-peve-MS2-LacZ* was also inserted in a different landing site in the third chromosome - *attP86Fb* (BDSC # 24749).

##### Expression of ectopic NICD

To generate *eve2-NICD* the plasmid 22FPE (Kosman and Small 1997), which contains 2 copies of the *eve2* enhancer with five high affinity *bicoid* sites, FRT sites flanking a transcription termination sequence and the *eve 3’UTR*, was transferred to *pGEM-t-easy* using *Eco*RI sites and from there to *pattB* (Bischof et al. 2013) using a *Not* I site. The NICD fragment from Notch was excised from an existing *pMT-NICD* plasmid and inserted in *pattB-22FPE* through the *Pme*I site to create the *pattB-eve2x2-peve-FRT-STOP-FRT-NICD-eve3’UTR* construct (referred to as *eve2-NICD*). This was inserted into the *attP* landing site at 51D in the second chromosome. To increase the amount of ectopic NICD produced, the same *eve2-NICD* construct was also inserted in the *attP40* landing site at 25C and recombined with *eve2-NICD* 51D to produce 2x*eve2-NICD*. Sequences of all generated plasmids are available in a benchling repository.

#### Fly strains and genetics

To observe the expression pattern and dynamics from *m5/m8-peve*, *sim-peve*, *m8NE-peve* and the different promoter combinations (Fig. 1, S1) females expressing His2av-RFP and MCP-GFP (BDSC #60340) in the maternal germline were crossed with males expressing the *MS2-lacZ* reporter constructs.

To test expression from *m5/m8-peve* in the *Dl* and *neur* mutant backgrounds, *His2Av-RFP* from *His2av-RFP; nos-MCP-GFP* (BDSC #60340) was recombined with *nos-MCP-GFP* in the second chromosome (BDSC #63821) and combined with a deficiency encompasing the *Dl* gene (*Df(3R)Dl^FX3^*, (Vässin and Campos-Ortega 1987)) or a *neuralized* loss of function allele (*neur*[*^11^*], BDSC #2747). *m5/m8-peve-MS2-lacZ* was also combined with the *Dl* and *neur* alleles and mutant embryos were obtained from the cross *His2Av-RFP,nos-MCP-GFP; mut / TTG* x *m5/m8-peve-MS2-lacZ; mut / TTG*. Homozygous mutant embryos for *Dl* or *neur* were selected by the lack of expression from the *TTG* balancer (*TM3-twi-GFP*, BDSC #6663).

To observe transcripion from two MS2 reporters in each cell (Fig. 2, S2) *His2Av-RFP* (BDSC #23650) was recombined with *nos-MCP-GFP* (from BDSC #60340) in the third chromosome and combined with *m5/m8-peve*, *sim-peve* or *m5/m8*^Δ*twi*^ ^Δ*dl*^*-peve* MS2 reporters. *m5/m8-peve* x2, *sim-peve* x2 and *m5/m8*^Δ*twi*^ ^Δ*dl*^*-peve* x2 embryos were obtained from the stocks *m5/m8-peve-MS2-LacZ; His2Av-RFP,nos-MCP-GFP*, *sim-peve-MS2-LacZ; His2Av-RFP,nos-MCP-GFP* and *m5/m8*^Δ*twi*^ ^Δ*dl*^*-peve-MS2-LacZ; His2Av-RFP,nos-MCP-GFP*, respectively; while *m5/m8-peve* + *sim-peve* embryos were obtained from crosssing *sim-peve-MS2-LacZ; His2Av-RFP,nos-MCP-GFP* females with *m5/m8-peve-MS2-LacZ* males.

To observe transcription from MS2 reporters in conditions of ectopic Notch activity the *FRT-STOP-FRT* cassette had to be first removed from the *eve2-NICD* construct by expression of a flippase in the germline. To do so flies containing *ovo-FLP* (BDSC #8727), *His2Av-RFP* and *nos-MCP-GFP* were crossed with others containing *eve2-FRT-STOP-FRT-NICD*, *His2Av-RFP* and *nos-MCP-GFP*. The offspring of this cross (*ovo-FLP/+; eve2-FRT-STOP-FRT-NICD/+; His2Av-RFP, nos-MCP-GFP*) induced FRT removal in the germline and were crossed with the MS2 reporters to obtain embryos expressing ectopic NICD. We note that only half of the embryos present the *eve2-NICD* chromosome, which could be distinguished by ectopic MS2 activity and an ectopic cell division of all the cells in the *eve2* stripe after gastrulation. The other 50% embryos obtained from this cross were used as the wild type controls. This strategy was used to observe transcription from *m5/m8-peve*, *sim-peve*, *m8NE-peve*, *m5/m8-pm5*, *sim^SPS^-peve*, *m5/m8^insSPS^-peve*, *m5/m8*^Δ*twi*^*-peve*, *m5/m8*^Δ*dl*^*-peve* and *m5/m8*^Δ*twi*^ ^Δ*dl*^*-peve*. To measure transcription from 2x*eve2-NICD* (Fig. 4, S4) removal of the *FRT-STOP-FRT* cassete was induced from the male germline to avoid recombination. To do so, *betaTub85D-FLP* (BDSC #7196) females were crossed with 2x*eve2-NICD* males and the male offspring of this cross (*betaTub85D-FLP/Y; 2xeve2-NICD* /+), which induces FRT removal in the germline, were crossed with *m5/m8-peve-MS2-lacZ; His2AvRFP, nos-MCP-GFP* or *sim-peve-MS2-lacZ; His2AvRFP, nos-MCP-GFP* females. As in the previous strategy, only half of the embryos presented the 2x*eve2-NICD* chromosome and were distinguished by the ectopic activity. To express two *m5/m8-peve* reporters in conditions of ectopic NICD activity, *m5/m8-peve* and *eve2-NICD* were recombined in the second chromosome and embryos were obtained by crossing *m5/m8-peve-MS2-lacZ; His2AvRFP, nos-MCP-GFP* females with *betaTub85D-FLP/Y; m5/m8-peve, eve2-NICD* /+ males. Embryos were selected by the presence of two MS2 spots in each cell, which also ectopically expressed NICD.

### METHOD DETAILS

#### Live imaging

Embryos were dechorionated in bleach and mounted in Voltalef medium (Samaro) between a semi-permeable membrane and a coverslip. The ventral side of the embryo was facing the coverslip in all movies except when looking at transcription in the DE region, for which they were mounted laterally. Movies were acquired in a Leica SP8 confocal using a 40x apochromatic 1.3 objective and the same settings for MCP-GFP detection: 40mW 488nm argon laser detected with a PMT detector, pinhole airy=4. Other settings were slightly different depending on the experiment. To observe transcription in the whole embryo (Fig. 1 and S1) settings were: 3% 561nm laser, 0.75x zoom, 800×400 pixels resolution (0.48um/pixel), 19 1um stacks, final temporal resolution of 10 seconds/frame). To observe transcription from 2 MS2 alleles simultaneously (Fig. 2, S2 and S6**E**) settings were: 2% 561nm laser, 1.5x zoom, 800×400 pixels resolution (0.24um/pixel), 29 1um stacks, final temporal resolution of 15s/frame). In all other experiments with ectopic NICD a ~150×150um window anterior to the center of the embryo was captured. Settings were: 2% 561nm laser, 2x zoom, 400×400 pixels resolution (0.36um/pixel), 29 1um stacks, final temporal resolution of 15s/frame). All images were collected at 400Hz scanning speed in 12 bits.

### QUANTIFICATION AND STATISTICAL ANALYSIS

#### Image analysis

Movies were analyzed using custom Matlab (Matlab R2018a, Mathworks) scripts (available at GitHub:FryEmbryo3DTracking. Briefly, the His2Av-RFP signal was used to segment and track the nuclei in 3D. Each 3D stack was first filtered using a median filter, increasing the contrast based on the profile of each frame to account for bleaching and a fourier transform log filter (Garcia et al. 2013). Segmentation was performed by applying a fixed intensity threshold, 3D watershed accounting for anisotropic voxel sizes (Mishchenko 2015) to split merged nuclei and thickening each segmented object. Nuclei were then tracked by finding the nearest object in the previous 2 frames which was closer than 6 um. If no object was found, that nuclei was kept with a new label, and only one new object was allowed to be tracked to an existing one. After tracking, the 3D shape of each nucleus in each frame was used to measure the maximum fluorescence value in the GFP channel, which was used as a proxy of the spot fluorescence. We note than when a spot cannot be detected by eye this method detects only background, but the signal:background ratio is high enough that the subsequent analysis allows to classify confidently when the maximum value is really representing a spot.

In experiments with two MS2 reporters the maximum intensity pixel per nucleus does not allow to separate transcription from the two alleles. To do so, the 3D Gaussian spot detection method from (Garcia et al. 2013) was implemented in the existing tracking, such that each spot was segmented independently and associated with the overlapping nuclei. In this manner only active transcription periods were detected and no further processing of the traces was required.

#### MS2 data processing

From the previous step we obtained the fluorescent trace of each nuclei over time. Only nuclei tracked for more than 10 frames were retained. First nuclei were classified as inactive or active. To do so the average of all nuclei (active and inactive) was calculated over time and fitted to a straight line. A median filter of 3 was applied to each nuclei over time to smooth the trace and ON periods were considered when fluorescent values were 1.2 times the baseline at each time point. This produced an initial segregation of active (nuclei ON for at least 5 frames) and inactive nuclei. These parameters were determined empirically on the basis that the filters retained nuclei with spots close to background levels and excluded false positives from bright background pixels. The mean fluorescence from MCP-GFP in the inactive nuclei was then used to define the background baseline and active nuclei were segregated again in the same manner. The final fluorescence values in the active nuclei were calculated by removing the fitted baseline from the maximum intensity value for each, and normalizing for the percentage that the MCP-GFP fluorescence in inactive nuclei decreased over time to account for the loss of fluorescence due to bleaching. Nuclei active in cycles before nc14 were discarded based on the timing of their activation.

In all movies, time into nc14 was considered from the end of the 13th syncythial division. When this was not captured the movies were synchronized by the gastrulation time. Plots showing mean fluorescent levels were obtained by calculating the mean and SEM of all fluorescent traces for multiple embryos aligned by the begining of nc14. Calculating the mean levels of multiple embryos taken individually returned very similar profiles, indicating there is little embryo-to-embryo variability. In figures 1**C** and 3**C** the total mRNA production per cell (in AU) was calculated by adding all the normalized fluorescent intensities for each nuclei.

Each embryo was classified into the 4 regions (ME, MSE, NE and DE) by drawing rectangular shapes in a single frame and finding which centroids overlapped with each region. In *eve2-NICD* these regions along the DV axis were defined within the *eve2* stripe (~ 6-7 cells wide in all movies). In wild type embryos ME and MSE regions were drawn in the whole field of view (~ 150×150 um anterior half of the embryo).

#### Definition of bursting properties

Bursts were defined as periods were the median-filtered signal was higher than 1.2 times the baseline for at least 5 frames within a period from 15 min into nc14. These defined the burst duration and the time off between bursts. The amplitude was defined as the mean value within each burst period. The proportion of active cells was defined as the percentage of cells that switch on at any point after 15 min in each of the defined regions. ‘Sustained’ transcription was defined as nuclei with at least one burst longer than 10 min. This was based on analyzing regions where separated burst of activity were detected (mesoderm and dorsal ectoderm) where most bursts were <10 min. Off periods shorter than circa 2 mins would not have been resolved because the MS2 loops were positioned within the 5’UTR and the limit of resolution depends on the time taken for a PolII molecule to complete transcription.

Onsets and ends of transcription were defined as the beginning of the first burst and the end of the last respectively (also starting at 15 min into nc14). In Figures 2 and S2 to be more precise in measuring the onsets and end-points of transcription for both MS2 alleles they were scored manually as the first and last frame a spot is detected and randomly assigned ‘allele 1’ or ‘allele 2’. The total variability was the variance of all onsets or end points, combining both alleles. The extrinsic variability was calculated as the covariance of onsets and ends between alleles 1 and 2. The remaining (total - covariance) corresponds to the intrinsic variability within each cell.

#### Statistical analysis

In figure legends, n number indicates number of embryos imaged for each biological condition. Where appropiate, n number next to heatmaps indicates total number of cells combining all embryos for each biological condition. Plots showing mean levels of transcription and SEM (standard error of the mean) combine all traces from multiple embryos from the same biological condition. Violin plots show the bursting properties (amplitude, burst duration and off period) for each independent burst in all traces in multiple embryos, therefore the n number can be significantly greater than the number of cells in each condition. Because these properties do not follow a normal distribution, their statistical significance was tested with two Kolmogorov-Smirnov test. Levels of significance are indicated in the figure legends.

### DATA AND SOFTWARE AVAILABILITY

Scripts for tracking and analysis of MS2 movies are available at GitHub:FryEmbryo3DTracking.

The code developed for the modelling approach to infer changes in parameters of transcription causing changes in mean levels of transcription is available at GitHub:FFR_ACF.

## KEY RESOURCES TABLE

## Supplemental Items

**Figure S1. Related to Figure 1. The temporal profile of transcription is characteristic of MSE enhancers.**

**Figure S2. Related to Figures 2 and 3. Quantification of the variability intrinsic and extrinsic to transcription.**

**Figure S3. Related to Figure 4. Modelling a two-state promoter to infer changes in the kinetic parameters of transcription.**

**Figure S4. Related to Figure 4. Effects of NICD on the transcriptional bursting properties.**

**Figure S5. Related to Figure 5. Disruption of a SPS site produces lower transcription levels but does not delay the onset of transcription.**

**Figure S6. Related to Figure 6. Effects of mutations in Twist or Dorsal motifs in the onset of transcription.**

**Table S1. Related to STAR Methods. Primers used to amplify enhancer and promoter sequences and to introduce mutations in the enhancers.**

**Supplemental Methods: Modelling changes in kinetic parameters of transcription**

**Movie S1 Related to Figure 1. Expression of *m5/m8-peve*.** Movie showing transcription from *m5/m8-peve* in the mesectoderm stripe during nc14. Also note earlier, Notch independent transcription in broad domains in nc10-13 and in few scattered cells during the first few minutes of nc14, followed by a long period (approximately 20 min) of inactivity before MSE stripe cells initiate transcription. Maximum intensity projection (19×1um stacks) of the MCP-GFP (grey in left pannel, green in right pannel) and His2Av-RFP (blue in right pannel) channels. 0.36 um/px XY resolution and final time resolution of 10s/frame. Anterior to the left; embryo imaged from the ventral side.

**Movie S2 Related to Figure 1. Expression of *sim-peve*.** Movie showing transcription from *sim-peve* in the mesectoderm stripe and some mesodermal cells during nc14. Also note activity in scattered cells before nc14. Maximum intensity projection (29×1um stacks) of the MCP-GFP (grey in left pannel, green in right pannel) and His2Av-RFP (blue in right pannel) channels. 0.36 um/px XY resolution and final time resolution of 15s/frame. Anterior to the left; embryo imaged from the ventral side.

**Movie S3 Related to Figure 3. Ectopic expression of *m5/m8* with *eve2-NICD*.** Movie showing ectopic transcription from *m5/m8-peve* in the *eve2* domain during nc14. Maximum intensity projection (29×1um stacks) of the MCP-GFP (grey in left pannel, green in right pannel) and His2Av-RFP (blue in right pannel) channels. 0.36 um/px XY resolution and final time resolution of 15s/frame. Anterior to the left; embryo imaged from the ventral side.

**Movie S4 Related to Figure 3. Ectopic expression of *sim* with *eve2-NICD*.** Movie showing ectopic transcription from *sim-peve* in the *eve2* domain during nc14. Maximum intensity projection (29×1um stacks) of the MCP-GFP (grey in left pannel, green in right pannel) and His2Av-RFP (blue in right pannel) channels. 0.36 um/px XY resolution and final time resolution of 15s/frame. Anterior to the left; embryo imaged from the ventral side.

**Movie S5 Related to Figure 3. Regions of ectopic expression of *m5/m8* and *sim* with *eve2-NICD*.** Combined movie of *m5/m8-peve* (left) and *sim-peve* (right) showing ectopic transcription in the *eve2* stripe. The maximum projection of the MCP-GFP signal is overlayed with tracked nuclei false colored with the maximum intensity pixel in each nuclei. Active nuclei in each of the analyzed regions is marked with a different color: red (mesoderm), purple (mesectoderm), blue (neuroectoderm) and green (dorsal ectoderm). Anterior to the left; embryo imaged from the ventral side.

**Movie S6 Related to Figure 3. Transcription of two *m5/m8-peve* MS2 reporters in the presence of ectopic NICD.** Movie showing ectopic transcription from *m5/m8-peve*x2 in the *eve2* domain during nc14. Maximum intensity projection (29×1um stacks) of the MCP-GFP (grey in left pannel, green in right pannel) and His2Av-RFP (blue in right pannel) channels. 0.36 um/px XY resolution and final time resolution of 15s/frame. Anterior to the left; embryo imaged from the ventral side.

