## Supplemental Figure and Methods for "Enhancer priming enables fast and sustained transcriptional responses to Notch signaling"

### Supplemental Figures

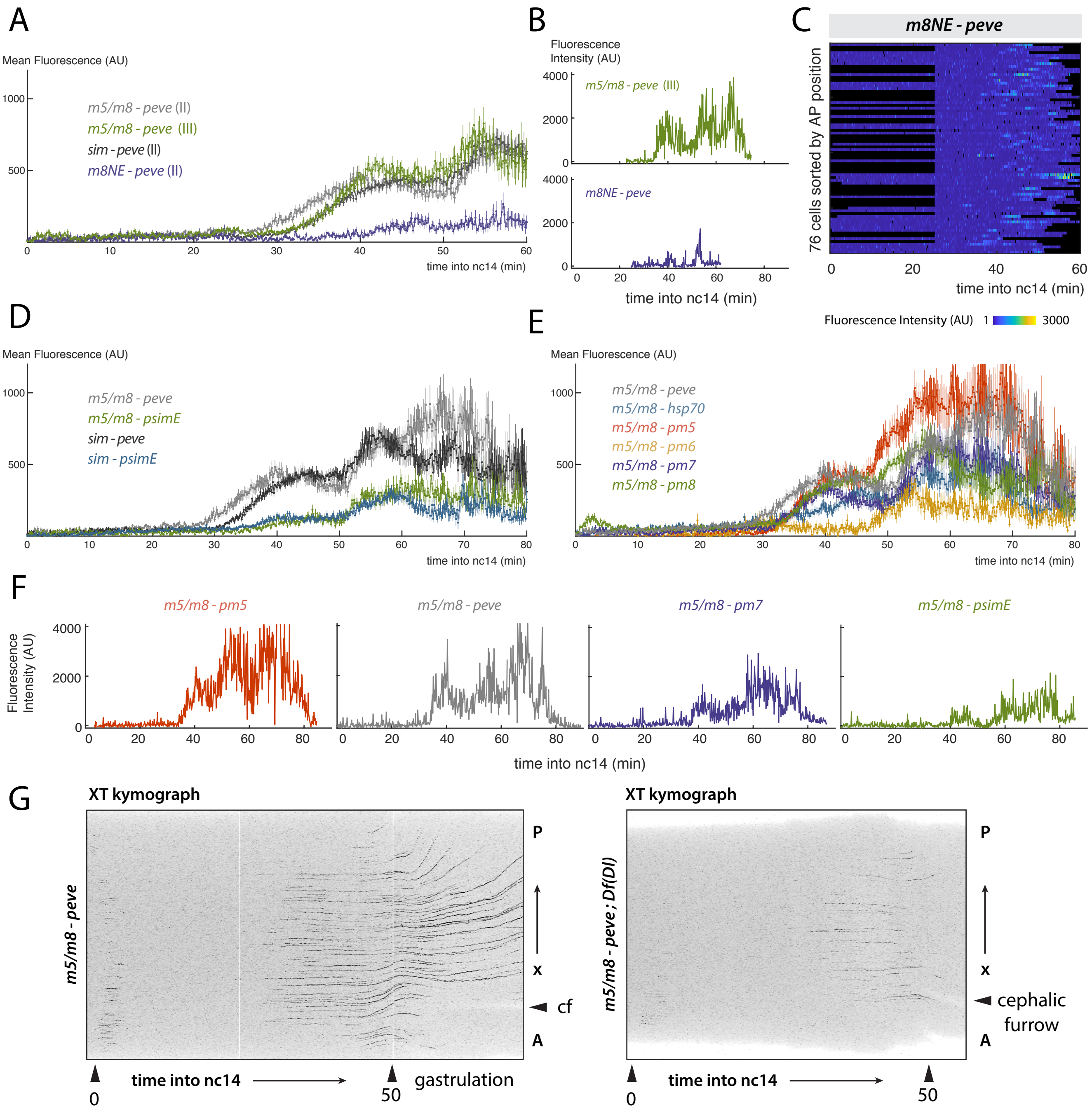


**Figure S1. Related to Figure 1. The temporal profile of transcription is characteristic of MSE enhancers. A**) Insertions of *m5/m8-peve* in landing sites in the second (light grey) and third (green) chromosome present the same temporal pattern and mean levels; and a Notch responsive neuroectodermal enhancer (*m8NE*, purple) presents a different temporal pattern than *m5/m8* and *sim*. **B**) Examples of traces from an *m5/m8-peve* insertion in a different genomic location, showing a continuous profile similar to Fig. 1D (top) and from *m8NE-peve*, a neuroectodemal enhancer that produces 'bursty' transcription in the MSE. **C**) *m8NE* produces asynchronized transcription in the MSE. **D**) The early promoter of *sim* (*psimE*) produces similar, lower mean levels of transcription from *m5/m8* and *sim* compared to the *eve* promoter. **E**) Different promoters from *E(spl)* complex genes and *hsp70* also affect the mean levels of activity but not the global pattern of transcription. **F**) Examples of fluorescent traces from different promoters. All produce continuous traces of different levels. **G**) Projections of the raw MCP-GFP channel over the Y and Z axes creating an XT kymograph. Only a few cells initiate transcription in embryos lacking zygotic Dl protein (right) compared to wild type embryos (left) and it is extinguished earlier. Mean and SEM are shown in **A**, **D** and **E**. Grey lines are re-plotted from Figs. 1D and 1G for comparison. $n = 3$ (*m5/m8-peve*III), 2 (*m8NE-peve*), 2 (*m5/m8-psimE*), 4 (*sim-psimE*), 3 (*m5/m8-hsp70*), 3 (*m5/m8-pm5*), 3 (*m5/m8-pm6*), 3 (*m5/m8-pm7*), 4 (*m5/m8-pm8*).

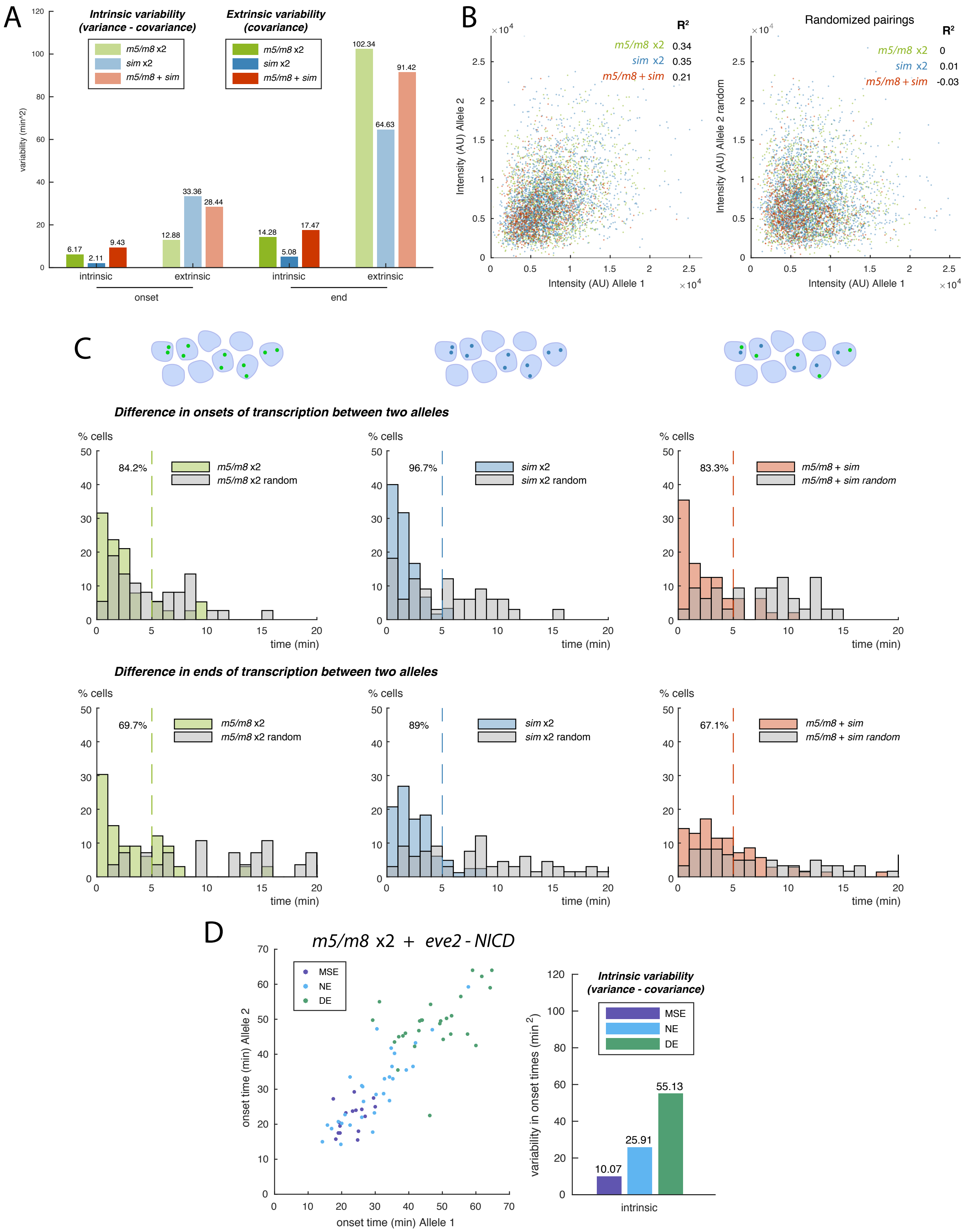


**Figure S2. Related to Figures 2 and 3. Quantification of the variability intrinsic and extrinsic to transcription. A**) Intrinsic (total variability minus covariance) and extrinsic (covariance) variability quantified in the onsets and ends of transcription using two MS2 reporters per cell. The amount of intrinsic variability is much smaller than the extrinsic and the intrinsic variability is higher in the ends than onsets of transcription for each combination. **B**) The fluorescence

**Figure S2 (continued).** intensities in two alleles at any timepoint present a small but significant correlation (left), compared to a correlation of 0 when the allele pairs are randomly assigned (right). Each color indicates the combination of 2 reporters compared. **C**) Histograms of the time difference between the appearance or dissapearance of transcription foci between the two reporters. The synchrony in the onset times is less than 5 min in more than 80% of the cells and more than 60% in the ends of transcription. Grey bars indicate the distribution of time differences when the allele pairs are randomly assigned. **D**) In conditions of ectopic Notch activity, nuclei in different regions present different intrinsic variability in the onset times of activation.

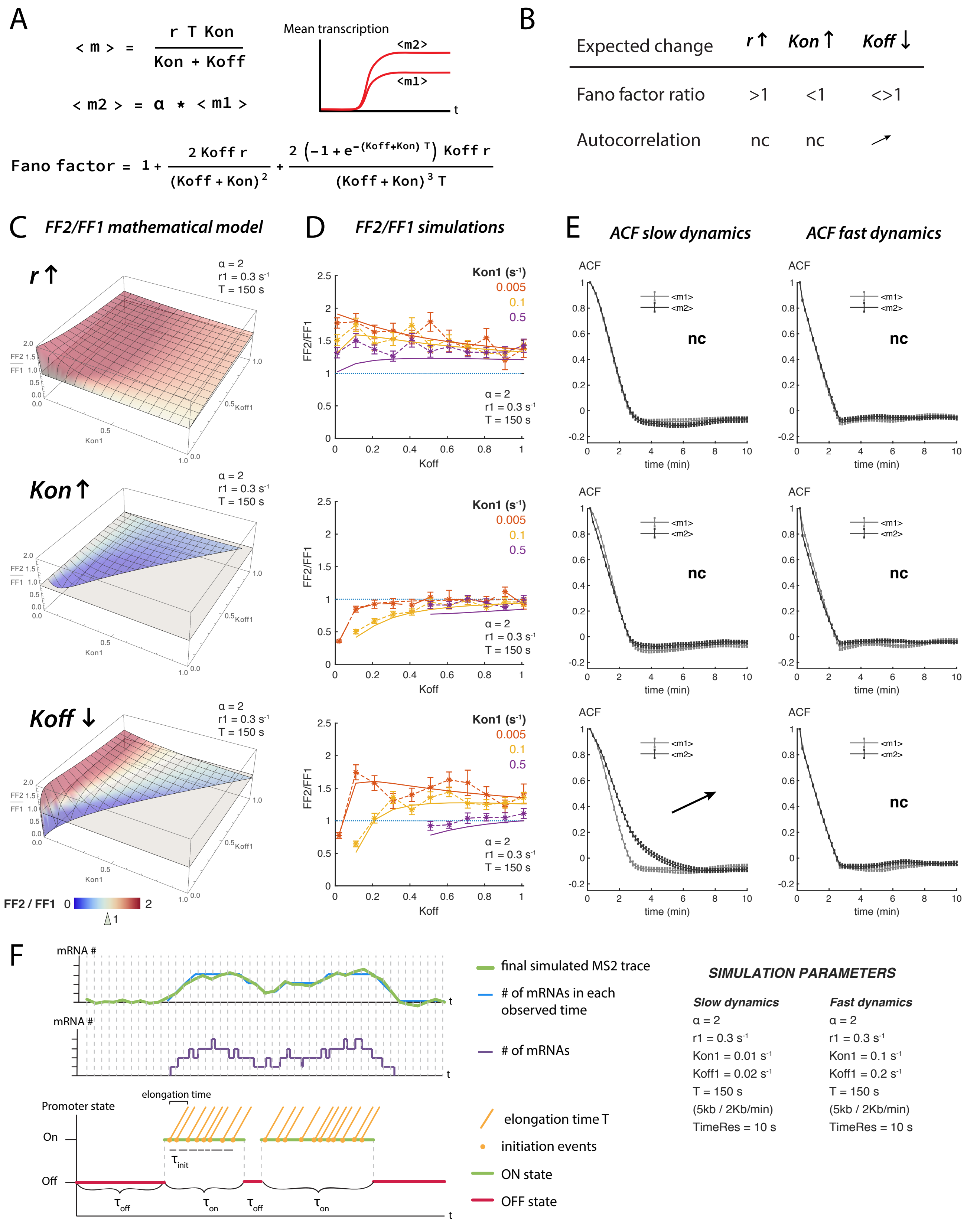


**Figure S3. Related to Figure 4. Modelling a two-state promoter to infer changes in the kinetic parameters of transcription.**

**Figure S3 (continued).** **E**) Expressions for the mean and Fano factor of the described 2-state model of transcription. Simulations and experiments compare the traces from two populations that have distinct means $\langle m1 \rangle$ and $\langle m2 \rangle$. $\alpha$ is the fold change in mean levels. The mean levels of transcription could increase from an increase in $r$, increase in $K_{on}$ or decrease in $K_{off}$. **F**) Summary of the effects that modifying each parameter to produce an increase of $\alpha$ in the mean have on the Fano factor ratio (FFR = FF2/FF1) and autocorrelation function (ACF). When $r$ increases, all FFR values are greater than 1 and no change (nc) in the ACF is observed. When $K_{on}$ increases all FFR values are smaller than 1 and no change is observed in the ACF. When $K_{off}$ decreases FFR values can be greater or smaller than 1 and the ACF presents a shift to the right when the dynamics are slow enough (see below). **G**) 3D plots representing the expected Fano factor ratio values from the mathematical model as a function of $K_{on1}$ and $K_{off1}$. $\alpha = 2$, $r_1 = 0.3s^{-1}$ and $T = 150s$ in the three plots. The grey surface indicates FFR = 1. Only $K_{on1}$ and $K_{off1}$ values that produce allowed (ie. positive) $K_{on2}$ and $K_{off2}$ values are plotted (see Supplementary Methods for details). Surface map is colored based on FFR values ranging from 0 (blue) to 2 (red). When an increase of $\alpha$ in the mean is caused by an increase in $r$ all FF ratio (FF2/FF1) values for any $K_{on}$ and $K_{off}$ values are greater than 1 (top plot). When it is due to an increase in $K_{on}$ all FF ratios are smaller than 1 (middle plot). When $K_{off}$ decreases to produce an increase of $\alpha$ in the mean, the obtained FF ratio values can be greater or smaller than 1 depending on the starting $K_{on1}$ and $K_{off1}$ parameters (bottom plot). **H**) Comparisons of the Fano factor ratios obtained from simulations of MS2 traces with different parameters (dashed lines) and the predicted from the mathematical model (solid line). Asterisks and error bars are mean and SD of the Fano factor ratio over 50 bootstraps of 1000 simulated MS2 traces, using the described $K_{on1}$ and $K_{off1}$ values and $\alpha = 2$, $r_1 = 0.3s^{-1}$, $T = 150s$ $(5Kb\ /\ 2Kb/min)$. The expected trends in Fano factor ratios are correctly recovered in the simulations of transcription. **I**) Plots showing the changes ACF over time in simulated traces, comparing mean and SD of the ACF of 200 simulated MS2 traces in 50 bootstraps obtained from two groups: $\langle m1 \rangle$, grey, and $\langle m2 \rangle$, black. The parameters used for the simulations are $K_{on1} = 0.01$ and $K_{off1} = 0.02$ (slow dynamics, left column) or $K_{on1} = 0.1$ and $K_{off1} = 0.2$ (fast dynamics, right column) and $\alpha = 2$, $r_1 = 0.3s^{-1}$, $T = 150s$. No changes in the ACF are observed when the dynamics are fast. When the dynamics are slow, increases in $r$ or $K_{on}$ do not produce any change in the ACF but changes decreases in $K_{off}$ shift the ACF to the right, from $\langle m1 \rangle$ to $\langle m2 \rangle$. **J**) Schematic representation of the steps to simulate MS2 traces. First ON and OFF states are generated based on the Gillespie algorithm, ON states are filled with initiation events that spread over their elongation time $T$. The final trace is obtained by counting the number of initiation events at each of the observed time points and adding gaussian noise to simulate experimental noise (see Supplementary Methods).

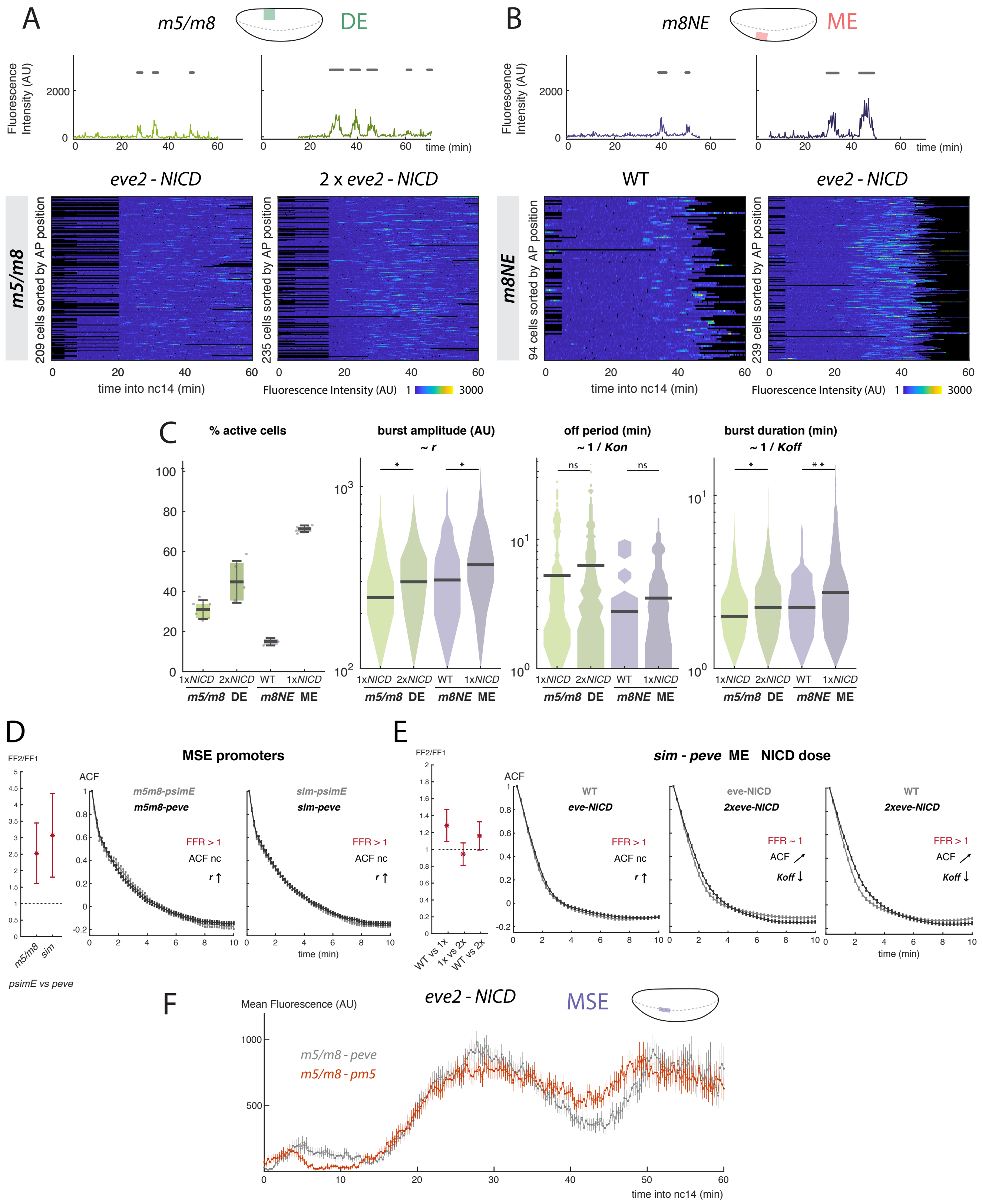


**Figure S4. Related to Figure 4. Effects of NICD on the transcriptional bursting properties. A**) Example traces and heatmaps of cells showing bursts of transcriptional activity from *m5/m8* in the dorsal ectoderm region in conditions of ectopic Notch activity. **B**) Example traces and heatmaps of cells showing bursts of transcriptional activity from *m8NE* in the mesoderm in wild type and *eve2-NICD* embryos. Burst periods are marked with a grey line. **C**) Quantification of the effects of NICD levels on the bursting properties. In both enhancers higher NICD produces a greater proportion of active cells and bigger bursts (increased amplitude and duration). **D**-**E**) Plots showing the Fano factor ratio and changes

**Figure S4 (continued).** in ACF over time (FFRatio in red, ACF in grey/black plots). FFRatio plots mean and SD of the FFRatio (FF2/FF1) in 50 bootstraps. Dashed line indicates 1 to compare the obtained FFRatio values. ACF plots compare mean and SD of the ACF of all available MS2 traces in 50 bootstraps from two conditions (grey and black lines as indicated, the mean levels are always higher in the condition plotted with a black line). **D**) Analysis of traces from reporters containing different promoters reveals changes in the mean are due to changes in $r$ (FFRatio greater than 1 and no changes in the ACF). **E**) Comparison of the FF ratio and ACF in ME traces from *sim* in WT, *eve2-NICD* and 2x*eve2-NICD* reveals changes in the mean are consistent with increases in $r$ (WT vs *eve2-NICD* comparison, left) or decreases in $K_{off}$ (middle and right plots comparing *eve2-NICD* vs 2x*eve2-NICD* and WT vs *eve2-NICD*; ACF shifts to the right from the lower to higher mean condition). Note that the model assumes only one parameter changes. **F**) Higher NICD levels saturate the response from the effect on the enhancer. A promoter that produces higher mean levels in wild type embryos does not increase the levels with *eve2-NICD*. Differential distributions in **C** tested with two-sample Kolmogorov-Smirnov test: pvalues $<0.01$(*), $<10^{-5}$(**), $<10^{-10}$(***). n = 6 (*m5/m8 eve2-NICD* lateral view), 5 (*m5/m8* 2x*eve2-NICD* lateral view), 3 (*m8NE* WT), 5 (*m8NE eve2-NICD*) and 5 (*m5/m8-pm5 eve2-NICD*) embryos. Grey lines in **F** are re-plotted from Fig. 3H.

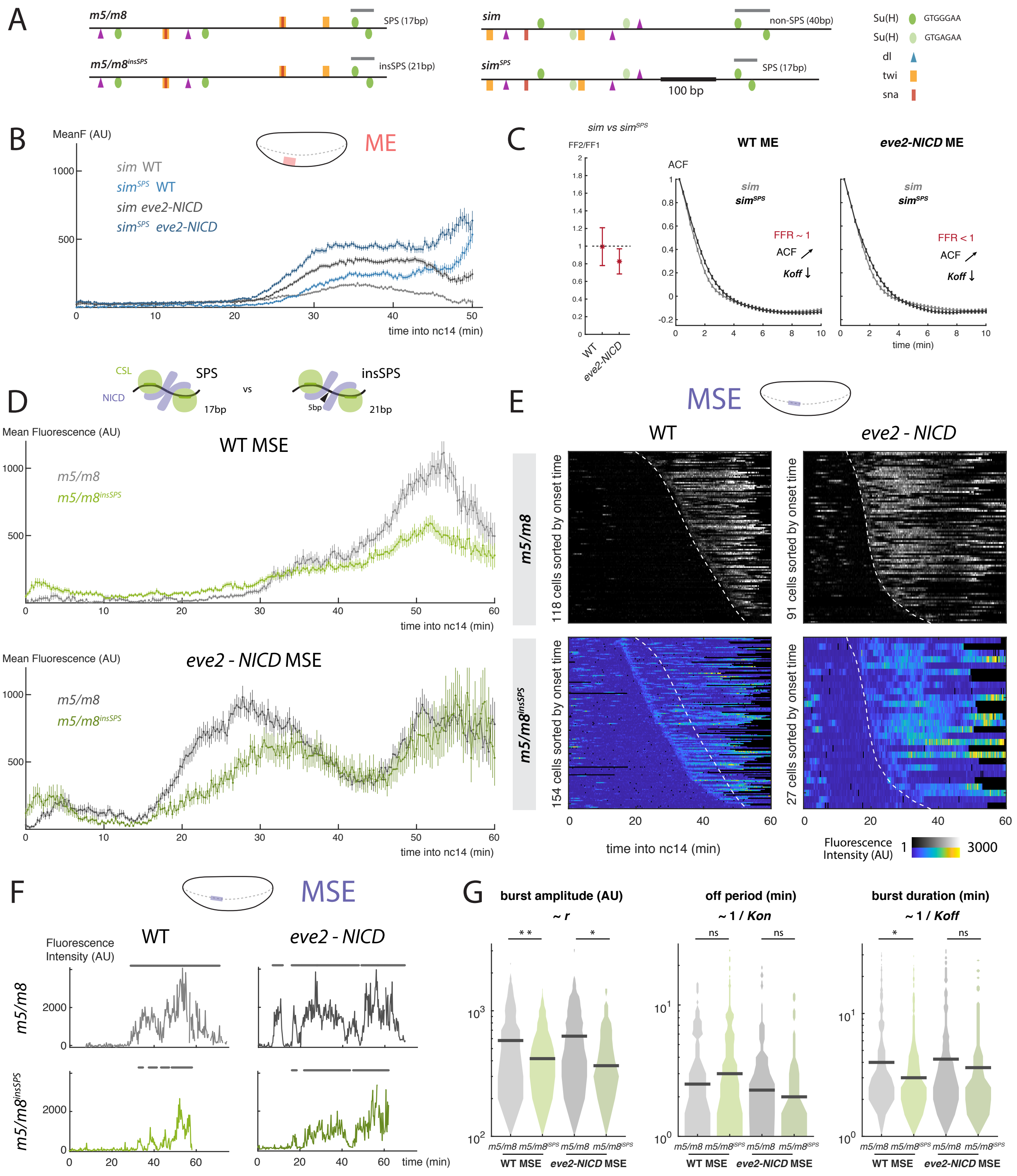

**Figure S5. Related to Figure 5. Disruption of a SPS site produces lower transcription levels but does not delay the onset of transcription. A**) Schematic representation of Su(H), Dorsal, Twist and Snail binding motifs in *m5/m8* and *sim* and introduced alterations in the SPS sites. **B**) $sim^{SPS}$ produces higher mean levels in the mesoderm compared to *sim*, in both wild type and *eve2-NICD* embryos. **C**) Plots showing the Fano factor ratio and changes ACF over time (FFRatio in red, ACF in grey/black plots). The Fano factor ratio and autocorrelation function of *sim* and $sim^{SPS}$ traces in the mesoderm in wild type and *eve2-NICD* embryos are compatible with changes in $Koff$ (shift in ACF) to produce increases in mean levels from *sim* to $sim^{SPS}$, in agreement with 5D. **D**) $m5/m8^{insSPS}$ produces lower mean levels of transcription compared to *m5/m8* but does not delay the onset of the response. **E**) $m5/m8^{ins}$ does not shift the onset of the response in *eve2-NICD* embryos (bottom) compared to *m5/m8* but presents some de-repression in wild type embryos (top). Dashed lines indicate onset times in the wild type enhancer. **F**) Examples of fluorescent traces in the mesectoderm region

**Figure S5 (continued).** in the described conditions. Burst periods are marked with a grey line. **G**) Quantification of the busting properties in the mesectoderm. $m5/m8^{insSPS}$ produces smaller bursts (lower amplitude and shorter duration) than *m5/m8*. Differential distributions in **G** tested with two-sample Kolmogorov-Smirnov test: pvalues $<0.01$(*), $<10^{-5}$(**), $<10^{-10}$(***). n = 5 ($m5/m8^{insSPS}$ WT), 3 ($m5/m8^{insSPS}$ *eve2-NICD*). Grey lines and heatmaps in **DE** are re-plotted from Fig. 3GH. **C** shows mean and SD over time of the mean Fano factor ratio and mean ACF over 50 bootstraps of all traces.

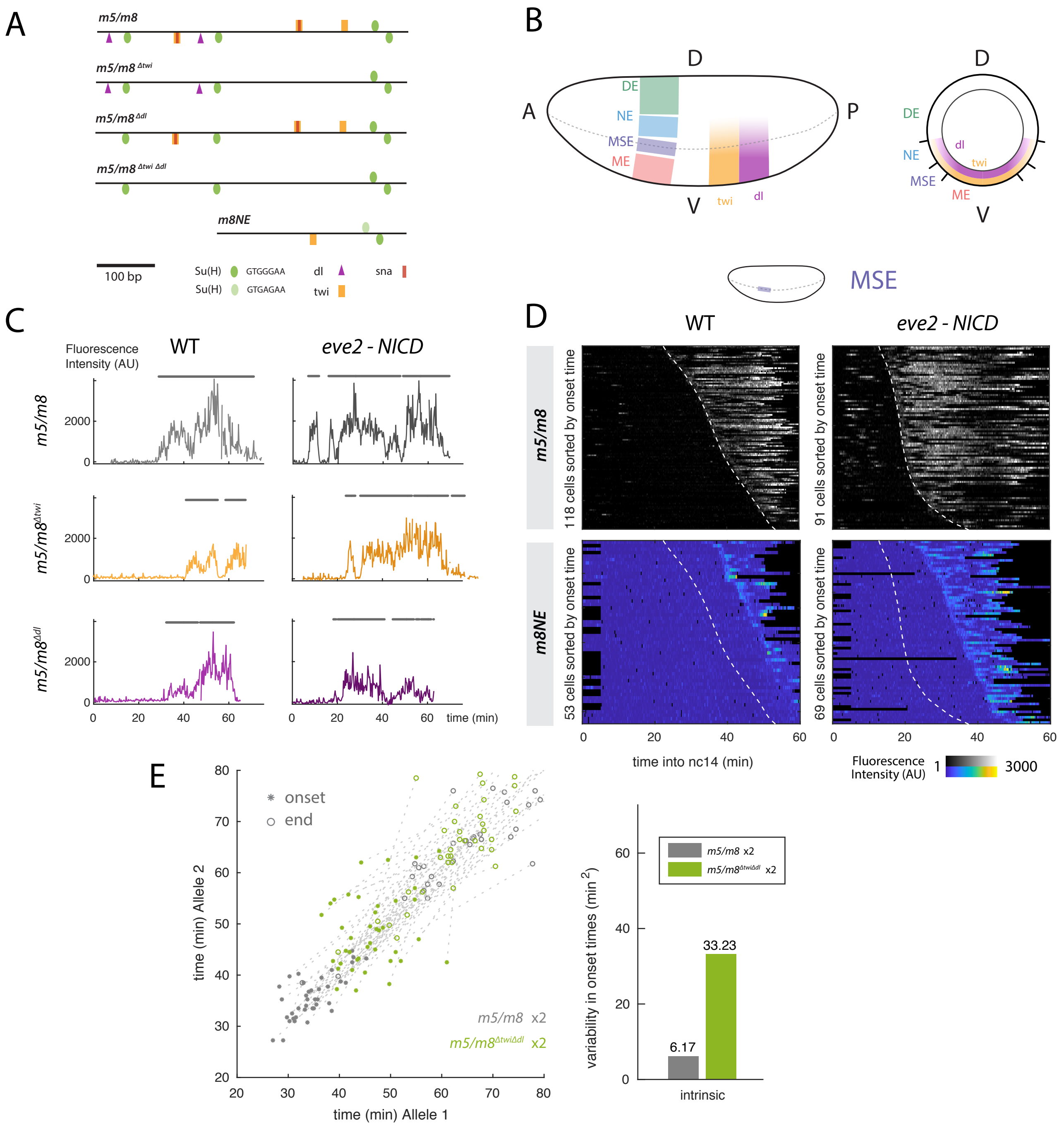


**Figure S6. Related to Figure 6. Effects of mutations in Twist or Dorsal motifs in the onset of transcription.** **A**) Schematic representation of the introduced mutations in *m5/m8* and comparison with a neuroectodermal enhancer, *m8NE*. **B**) Diagram of Twist and Dorsal gradients in the blastoderm embryo, showing lateral view (left) and cross-section (right). Both gradients extend in a ventral to dorsal gradient in the ME, MSE and NE. **C**) Examples of transcription traces from mesectodermal cells expressing *m5/m8* with mutated Twist or Dorsal motifs. The onset of transcription is delayed but transcription still occurs in a sustained manner. **D**) Heatmaps of MSE cells expressing *m8NE*. The onset of transcription is delayed compared to *m5/m8*. Dashed lines indicate onset times in the *m5/m8*. **E**) Quantification of the intrinsic variability in the transcription from a *m5/m8* enhancer with mutated Twist and Dorsal sites. Onset and end times for two $m5/m8^{\Delta twi \Delta dl}$*-peve* reporters in the same cell are shown and compared to *m5/m8-peve* (left). The intrinsic variability in the onset times of $m5/m8^{\Delta twi \Delta dl}$*-peve* increases compared to *m5/m8*. Grey dots and bar are re-plotted from Figures 2B and S2**A** for comparison. Greyscale heatmaps are duplicated from Fig. 3G.

**Table S1. Related to STAR Methods. Primers used to amplify enhancer and promoter sequences and to introduce mutations in the enhancers**. Restriction sites for *Hind*III, *Age*I and *Eag*I are underlined.

| Primer name | Sequence |
|---|---|
| m5/m8 S | AAGCTTTGTTCCGTTTGGTAAAACCC |
| m5/m8 AS | ACCGGTCTTTCCACTGACATTCGAATC |
| sim S | AAGCTTCCCCGGCATATGTTACGCAC |
| sim AS | ACCGGTGGTTACAGGCAAACAGCAAAC |
| m8NE S | AAGCTTGGATCCCCTGCCCCTGCTC |
| m8NE AS | ACCGGTAACTTCGTAGGACGGAGGAC |
| peve S | AATGTCAGTGGAAAGACCGGTTTGCCTGCAGAGCGCAGCG |
| peve AS | TCCAAGGGCGAATTCACCGGCCGAACGAAGGCAGTTAGTTGTTGACTGT |
| hsp70 S | AATGTCAGTGGAAAGACCGGTGAGCGCCGGAGTATAAATAGA |
| hsp70 AS | TCCAAGGGCGAATTCACCGGCCGTATTCAGAGTTCTCTTCTTGTATTC |
| pm5 S | AATGTCAGTGGAAAGACCGGTACGCACGCACAGCATAGCAAT |
| pm5 AS | TCCAAGGGCGAATTCACCGGCCGAAGATTTGTAGAAATGTGCTGAGCTG |
| pm6 S | AATGTCAGTGGAAAGACCGGTTGGGATGATGTTGCTGCTG |
| pm6 AS | TCCAAGGGCGAATTCACCGGCCGTGTAGTATCACTTTACAGATAAGAGT |
| pm7 S | AATGTCAGTGGAAAGACCGGTAGTTTGCTCCGCAGGTGGT |
| pm7 AS | TCCAAGGGCGAATTCACCGGCCGATCTTTCGGAGGAGGTTATCCTG |
| pm8 S | AATGTCAGTGGAAAGACCGGTGCAGCTGTTCCTTGTGAAAAA |
| pm8 AS | TCCAAGGGCGAATTCACCGGCCGTTTGAAAAATTTTGTATTCGGCT |
| psimE S | AATGTCAGTGGAAAGACCGGTGTGTGAGTGTGGTGCATATAAATTTCGC |
| psimE AS | TCCAAGGGCGAATTCACCGGCCGGCGCACTCGCCGATGGTTAGTCA |
| sim for simSPS S | AAGTGTTTCCCACGATTCTGTCCTCCTTATGTGAAACTC |
| sim for simSPS AS | TCAAGTTTCCCACAAGATGGAAAGTGGAGAGTCCATAA |
| SPS from m5/m8 S | ATGGACTCTCCACTTTCCATCTTGTGGGAAACTTGAGG |
| SPS from m5/m8 AS | TTTCACATAAGGAGGACAGAATCGTGGGAAACACTTT |
| insSPS S | TGAGGGCAAAGAGGGGTGTTTCCCACGATTCGAAT |
| insSPS AS | TGGGAAACACCCCTCTTTGCCCTCAAGTTTCCCAC |
| mut Twi 1 S | ACTGATTTCCGTCCCAATGAGTCCCAAAATTGCACACATC |
| mut Twi 1 AS | TTTGGGACTCATTGGGACGGAAATCAGTATCTTACGGATT |
| mut Twi 2 S | CAAAATTCCCATTAGGACATCATCGGTTTGGCCCACTGTG |
| mut Twi 2 AS | AACCGATGATGTCCTAATGGGAATTTTGAGGGTGCCTTGC |
| mut Twi 3 S | CGGGACTCGCATTCGGACAACCTCCGATTATAACTTATAA |
| mut Twi 3 AS | ATCGGAGGTTGTCCGAATGCGAGTCCCGAGTCCGAGCTCC |
| mut dl 1 S | CCGTTTGGTGAGATCTCAAAAATCACATTCGAAAA |
| mut dl 1 AS | TGATTTTTGAGATCTCACCAAACGGAACAAAGCTT |
| mut dl 2 S | TCGCCTTGGGAGATCTCATTTCCGACATCCCAAAA |
| mut dl 2 AS | TCGGAAATGAGATCTCCCAAGGCGAAGATGTGTGC |

#### Supplemental Methods: Modelling changes in kinetic parameters of transcription

We used a two-state promoter model of transcriptional activation in which the promoter switches between OFF and ON with constants $K_{on}$ and $K_{off}$ and releases mRNAs at a rate $r$ when the promoter is ON (Fig. 4E). This model also accounts for the residence time of polymerase on DNA while transcribing the gene (the elongation time $T$), so it is capturing what the MS2 system detects, ie. the number of nascent mRNA on the gene, rather than overall levels of mRNA in the cell. We take as a starting point expressions from (Choubey et al. 2015) for the mean and variance of the number of nascent mRNAs ($m$) in steady state:

$$\langle m \rangle = \frac{rTK_{on}}{K_{on} + K_{off}} \tag{1}$$

$$Var(m) = \langle m \rangle \left[ 1 + \frac{2rK_{off}}{(K_{on} + K_{off})^2} + \frac{2rK_{off}}{(K_{on} + K_{off})^3} \left( \frac{e^{-T(Kon+Koff)} - 1}{T} \right) \right] \tag{2}$$

We take the elongation time, $T$, to be fixed for a given gene. Thus, according to equation 1, the levels of transcription could increase in three ways: by increasing $r$, increasing $K_{on}$, or decreasing $K_{off}$.

Thus, because of this degeneracy, observing a change in $\langle m \rangle$ is alone insufficient to determine which underlying bursting parameter is being tuned to drive that change. However, we can make progress by incorporating the intrinsic noise of transcription into our analysis, since equation 2 indicates that changes to bursting parameters that have equivalent effects on the mean may nonetheless lead to different noise signatures. To do this, we calculate the Fano factor, which is defined as the variance divided by the mean:

$$Fano(m) = \frac{Var(m)}{\langle m \rangle} \tag{3}$$

$$= 1 + \frac{2rK_{off}}{(K_{on} + K_{off})^2} + \frac{2rK_{off}}{(K_{on} + K_{off})^3} \left( \frac{e^{-T(Kon+Koff)} - 1}{T} \right) \tag{4}$$

Where we see that the expression for the Fano factor is identical to the quantity inside the brackets in equation 2.

Next, we examine how changes to each bursting parameter in turn will affect the Fano factor and Mean, respectively, demonstrating how these signatures can be used to uncover the drivers of observed changes between different experimental conditions.

##### Pol II Initiation Rate ($r$)

We start by considering the case when $r$ is modulated. In the discussions that follow, we assume a situation in which we are comparing two experimental conditions that exhibit observable differences in their mean rate of expression, $\langle m \rangle$:

$$\alpha\langle m_1 \rangle = \langle m_2 \rangle \tag{5}$$

Our goal is to determine whether the modulation of specific parameters corresponds reliably with changes in the mean and Fano factor. To do this, we undertake analysis of the functional form of the partial derivatives of these empirical measures with respect to each parameter.

From equation 1, we have:

$$\frac{\partial\langle m \rangle}{\partial r} = \frac{TK_{on}}{\kappa} \tag{6}$$

$$\frac{\partial\langle m \rangle}{\partial r} > 0 \tag{7}$$

Where, for convenience, we have introduced the shorthand $\kappa = K_{on} + K_{off}$. So we see that $\langle m \rangle$ is monotonic with $r$: an increase in $r$ always leads to an increase in the mean (and vice versa). The strict inequality applies because the right-hand-side of eq. 6 can be zeros *if* no expression occurs. For the fano Factor, we have:

$$\frac{\partial Fano}{\partial r} = \frac{2K_{off}}{\kappa^2}\left(1 + \frac{e^{-\kappa T} - 1}{\kappa T}\right) \tag{8}$$

$$\frac{\partial Fano}{\partial r} \geq 0 \tag{9}$$

Unlike the mean, it is possible that a change in $r$ could lead to *no* observable modulation in the Fano factor; however, this only holds for exceptionally small values of $\kappa T$. More importantly, we see that it is impossible for the Fano factor to decrease when $r$ is increased. Thus, we conclude that an increase in $r$ must coincide with an increase in both the mean rate of expression and in the Fano factor, ie. the ratio between the Fano factors $Fano(m_2)$ and $Fano(m_1)$ where $\langle m_2 \rangle = \alpha\langle m_1 \rangle$ would always be greater than 1 (Fig. S3**H**, top panel).

**Activation Rate ($K_{on}$)**

As with $r$, we begin by examining how $\langle m \rangle$ changes in response to a change in $K_{on}$:

$$\frac{\partial \langle m \rangle}{\partial K_{on}} = \frac{rT}{\kappa} - \frac{rTK_{on}}{\kappa^2} \tag{10}$$

$$= \frac{rT}{\kappa}(1 - \frac{K_{on}}{\kappa}) \tag{11}$$

$$\frac{\partial \langle m \rangle}{\partial K_{on}} \geq 0 \tag{12}$$

Thus, as with $r$, the mean rate of expression increases monotonically in response to increases in $K_{on}$. Next, for the Fano factor, we have:

$$\frac{\partial Fano}{\partial k_{on}} = 2rK_{off}\Big( - \kappa^{-3}(2 + e^{-\kappa T}) + \frac{3\kappa^{-4}}{T}(1 - e^{-\kappa T})\Big) \tag{13}$$

$$= -\frac{2rK_{off}}{\kappa^3}\Big(2 + e^{-\kappa T} - \frac{3(1 - e^{-\kappa T})}{\kappa T}\Big) \tag{14}$$

To gain further insight, we need to examine limiting cases for the quantity $\kappa T$, which encodes the relative magnitude of the elongation time and switching rates, and which dictates the noise characteristics of the system. We start with the case where $\kappa T << 1$:

$$\frac{\partial Fano}{\partial k_{on}} \approx -\frac{2rK_{off}}{\kappa^3}\Big(2 + 1 - \kappa T - \frac{3(1 + \kappa T - 1)}{\kappa T}\Big) \tag{15}$$

$$\approx -\frac{2rK_{off}}{\kappa^3}\Big(3 - \kappa T - 3\Big) \tag{16}$$

$$\approx -\frac{2rK_{off}}{\kappa^3}(0) \tag{17}$$

$$\approx -\frac{2rK_{off}}{\kappa^3}\Big(3 - \kappa T - 3\Big) \tag{18}$$

$$\frac{\partial Fano}{\partial k_{on}} \approx 0 \tag{19}$$

For the opposite limit, where $\kappa T >> 1$, we have:

$$\frac{\partial Fano}{\partial k_{on}} \approx -\frac{2rK_{off}}{\kappa^3}\Big(2 + 0 - \frac{3(1 - 0)}{\kappa T}\Big) \tag{20}$$

$$\approx -\frac{4rK_{off}}{\kappa^3} \tag{21}$$

$$\frac{\partial Fano}{\partial k_{on}} \leq 0 \tag{22}$$

So we see that, an increase in $\langle m \rangle$ that is driven by an increase in $K_{on}$ will coincide with a *decrease* in the Fano factor. Thus, unlike $r$, where the signs of the change in the mean and Fano factor are the same, we find that the signs of the changes in the mean and Fano factor are opposite in the case of changes driven by $K_{on}$, ie. the ratio between the Fano factors $Fano(m_2)$ and $Fano(m_1)$ where $\langle m_2 \rangle = \alpha \langle m_1 \rangle$ would always be smaller than 1 (Fig. S3**H**, middle panel).

**Off Rate ($K_{off}$)**

For the mean, we have:

$$\frac{\partial \langle m \rangle}{\partial K_{off}} = -\frac{rK_{on}}{\kappa^2} \tag{23}$$

$$\frac{\partial \langle m \rangle}{\partial K_{off}} \leq 0 \tag{24}$$

Thus, as expected, an increase in $K_{off}$ leads to a *decrease* in $\langle m \rangle$. In keeping with our treatment in the case of $K_{on}$, we next examine the functional form of the Fano factor in the small and large $\kappa T$ limits. For $\kappa T << 1$, we expand about $\kappa T = 0$ to obtain an expression for the Fano factor :

$$Fano \approx 1 + \frac{2rK_{off}}{(K_{on}+K_{off})^2} + \frac{2rK_{off}}{(K_{on}+K_{off})^3}\Big(\frac{1-\kappa T-1}{T}\Big) \tag{25}$$

$$\approx 1 \tag{26}$$

Thus, consistent with our findings for $K_{on}$ the Fano Factor is largely insensitive to changes in $K_{off}$ for small $\kappa T$. This holds for $r$ as well, though we did not state so explicitly above. Next, we approximate the large $\kappa T$ limit by setting $e^{-kT} = 0$:

$$Fano \approx 1 + \frac{2rK_{off}}{\kappa^2} + \frac{2rK_{off}}{\kappa^3}\Big(\frac{0-1}{T}\Big) \tag{27}$$

$$\approx 1 + 2r\Big(\frac{K_{off}}{\kappa^2} - \frac{K_{off}}{\kappa^2}\frac{1}{\kappa T}\Big) \tag{28}$$

$$\approx 1 + 2r\Big(\frac{K_{off}}{\kappa^2}\Big) \tag{29}$$

Differentiating, we obtain:

$$\frac{\partial Fano}{\partial_{off}} \approx 2r\Big(\frac{1}{\kappa^2} - \frac{2K_{off}}{\kappa^3}\Big) \tag{30}$$

$$\approx \frac{2r}{\kappa^2}\Big(1 - \frac{2K_{off}}{\kappa}\Big) \tag{31}$$

The expression above reveals that, unlike $r$ and $K_{on}$, the direction of the change of the Fano Factor in response to a change in $K_{off}$ not fixed, but depends upon the relative sizes of $K_{on}$ and $K_{off}$, ie. the ratio between the Fano factors $Fano(m_2)$ and $Fano(m_1)$ where $\langle m_2 \rangle = \alpha \langle m_1 \rangle$ could be smaller or greater than 1 (Fig.S3**H**, bottom panel). Numerical simulations confirm this result.

**Stochastic simulations**

We next tested with simulations whether the Fano factor ratio can be used as a diagnostic tool of the underlying changes in the mean. We used stochastic simulations of transcription based on the Gillespie algorithm (Gillespie 1976) of the same two-state promoter model but using additional parameters to more resemble the biological MS2 data (accounting for the time MS2 loops are detected, acquisition time and adding experimental noise, Fig. S3**J**).

We then tested whether we could recover the same trends in Fano factor ratios in the simulation as expected from the mathematical model. Indeed, using a variety of starting parameters we could recover similar Fano factor values as expected from the mathematical model (Fig. S3**H**). However, given that changes in $K_{off}$ can produce Fano factor ratios greater or smaller than 1, calculation of the Fano factor and comparing whether it is greater or smaller than 1 alone is not sufficient to infer which parameter is being modified to produce the observed changes in the mean.

**Utilizing the Autocorrelation Function (ACF)**

The results of our analysis thus far indicate that modulations in $r$ and $K_{on}$ lead to distinct, well defined signatures in mean and Fano factors of experimentally observed expression levels. However, the degeneracy of the Fano factor shift with respect to changes in $K_{off}$ necessitates the incorporation of an additional observable, if we are to be able to distinguish the underlying drivers of changes between experimental conditions. To this end, we utilize the empirical Autocorrelation Function of our experimental MS2 traces.

The ACF function provides information about the speed of the system and the elongation rate (Desponds et al. 2016; Lammers et al. 2018). Intuitively, the more rapid the time scale with which the system switches between activity states (the larger $\kappa$ is), the faster the ACF decays. We used the same simulations to test if the autocorrelation function changes in different ways depending on the modified parameters, to help distinguishing between the 3 scenarios to increase the mean. If the dynamics are fast (Fig. S3**I**, right column, $K_{on1} = 0.1s^{-1}$ and $K_{on1} = 0.2s^{-1}$) no changes in the ACF were observed in any of the three cases. When the dynamics are slower (Fig. S3**I**, left column, $K_{on1} = 0.01s^{-1}$ and $K_{on1} = 0.02s^{-1}$), then the AC function shifts to the right (from $\langle m_1 \rangle$ to

$\langle m_2 \rangle$) when $K_{off}$ decreases. No changes are observed when $r$ or $K_{on}$ increase.

Therefore looking at both the Fano factor ratio and the autocorrelation function (when the dynamics are slow enough), provides enough information to distinguish between the three ways in which the mean can change (Fig. S3**F**):

- increase in $r$: FFRatio $> 1$ and no change in ACF
- increase in $K_{on}$: FFRatio $< 1$ and no change in ACF
- decrease in $K_{off}$: FFRatio $< 1$ or $> 1$ and shift to the right in ACF

**Estimating Fano factor from empirical data**

When applied to real MS2 traces, raw fluorescence profiles from each cell were processed by applying a median filter of 3, removing the background baseline and normalizing for bleaching as described in the Methods section. When the onset of transcription was different between experiments (eg. WT vs *eve2-NICD*) they were shifted to compare equivalent times. The Fano factor was calculated as the intrinsic variability divided by the mean over time:

$$Fano = \frac{\sigma_i^2}{\langle m \rangle} \tag{32}$$

$$= \frac{Var(m) - CoVar(m)}{\langle m \rangle} \tag{33}$$

The intrinsic component was calculated by subtracting an estimation of the extrinsic variability form the total noise. The contribution from the extrinsic noise, normally calculated from the covariance of two transcription traces from the same cell, was calculated by using neighbouring nuclei as proxi of two loci in the same cell and calculating their covariance. Using the experiments where two MS2 reporters are present in each cell we validated the contribution from extrinsic noise is equivalent within cell and across neighbouring cells. Both FFRatio and ACF were calculated by doing 50 bootstraps of all available traces and calculating the mean and SD.

The code used to simulate MS2 traces and calculate the Fano factor and ACF is available at GitHub:FFR_ACF.
